# High-dimensional hippocampus representation facilitates learning of low-dimensional abstract representation and supports generalization

**DOI:** 10.64898/2026.10.24.761577

**Authors:** Bin Wang, W. Jeffrey Johnston, Stefano Fusi

## Abstract

Semantic knowledge encompasses facts, concepts, and their relationships. Recent work has proposed that some forms of semantic knowledge could be encoded in low-dimensional neural representations, such as the abstract, disentangled representational geometries observed in many cognitive brain areas. Experimental work has shown that hippocampal (HPC) lesions impair the formation of flexible and generalizable semantic knowledge. However, HPC representations are often high-dimensional and sparse. How can these representations contribute to the formation of low-dimensional abstract representation across these cognitive cortical areas? Here, we reconcile these observations using multiregional neural network models. Modeling HPC as a sparse, decorrelating autoencoder, we show that its high-dimensional representations promote the robust and efficient formation of low-dimensional abstract representations in downstream regions such as prefrontal cortex. This effect is most pronounced when the geometry of sensory inputs is misaligned with the geometry of desired outputs, requiring a more complex input-output transformation. Mathematical analysis shows that hippocampal expansion recoding suppresses misleading low-dimensional input structure and reshapes the loss landscape of downstream networks, allowing abstract semantic geometry to emerge reliably. The resulting abstract representations provide an inductive bias for sample-efficient learning of novel semantic-to-action mappings. Finally, we show that consolidating hippocampus-dependent representations into a hippocampus-independent pathway improves out-of-distribution generalization based on input structure. Together, these results suggest a circuit mechanism by which high-dimensional hippocampal representations can support, rather than oppose, the emergence of low-dimensional abstract representations.

## Introduction

To survive in an ever-changing environment, the brain must extract structure from experience and generalize it to novel scenarios. Such generalization is possible because the world contains regularities, described by facts, concepts, and their relationships, collectively referred to as semantic knowledge. As a concrete example of semantic knowledge, consider four visual images depicting either broccoli or a fox, each shown in green or orange (Fig. 1A). Each image can be associated with two binary semantic features: whether the object is a plant or an animal, and whether it is typically or rarely encountered in everyday life. Importantly, similarity in semantic space does not necessarily imply visual similarity in pixel space. Two images that share a semantic feature (e.g., both orange broccoli and green fox are “rarely seen”) may look very different at the pixel level, whereas visually similar images (e.g., the two green objects) may differ substantially in their semantic features. Semantic knowledge can be generally encoded in the structure of inputs, outputs, and their relationships, and effectively learning this semantic structure can support flexible downstream behavior. Schematically, this process is illustrated in Figure 1B.

**Figure 1:**
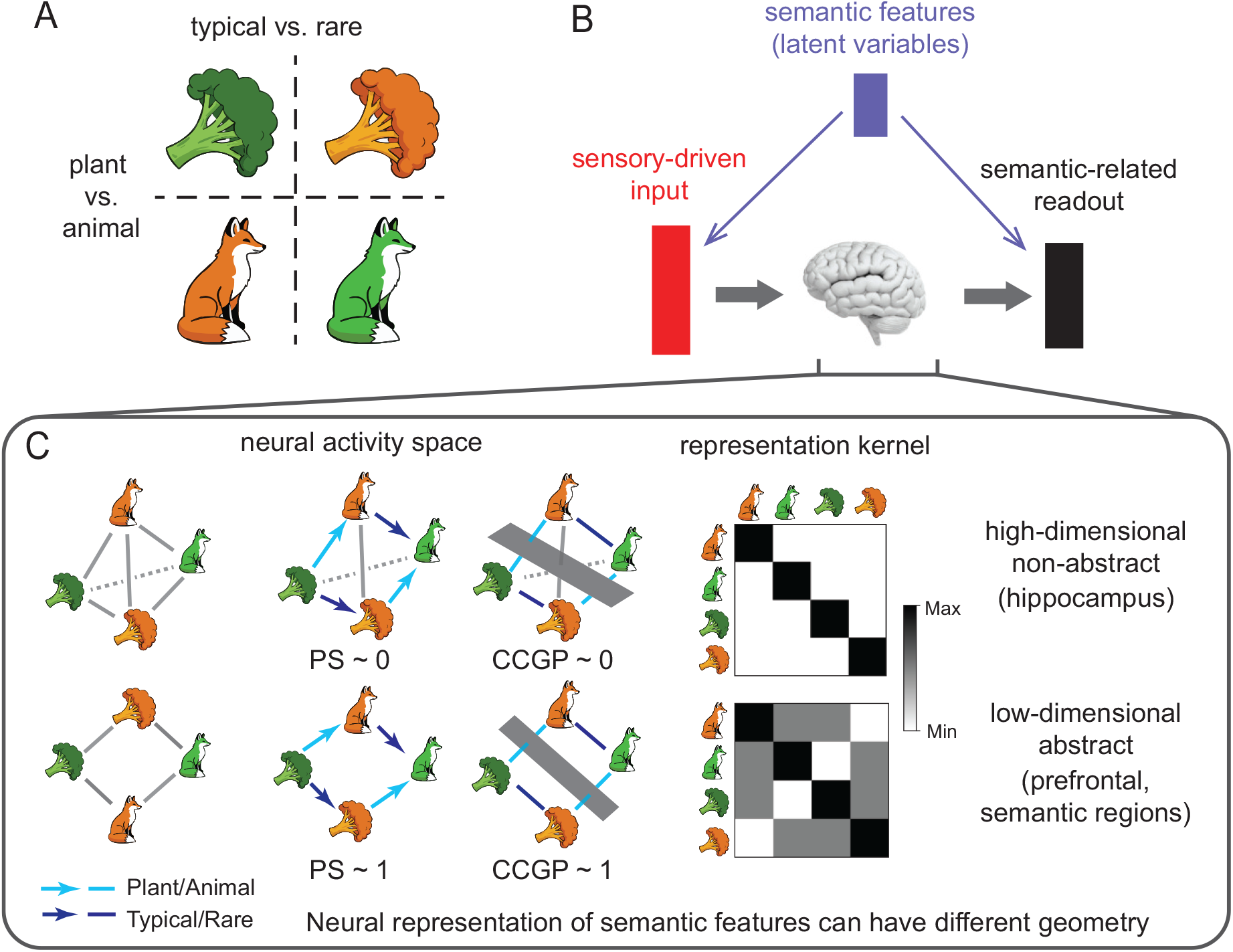
Semantically structured tasks and representational geometries of semantic variables. (A) Example set of four stimuli defined by two binary semantic features: animal or plant, and typical or rare. (B) Schematic of the semantically structured task, in which sensory-driven inputs are transformed into semantically related readouts, both determined by latent semantic features. (C) The same semantic variables can be encoded by different neural representational geometries. In a high-dimensional, non-abstract code (top), activity patterns form an approximately tetrahedral geometry, yielding a low parallelism score (PS) and a low cross-condition generalization performance (CCGP). In a low-dimensional, abstract code (bottom), each semantic variable is represented along a direction that remains invariant to changes in the other semantic variables, producing high PS and high CCGP. The corresponding representation kernels are shown on the right.

Semantic knowledge can be encoded in different forms in the brain. In particular, different brain regions may represent the same knowledge with distinct representation geometries at the neural population level. For the four stimuli above, there are multiple possible ways to encode both the stimuli and their semantic features (Fig. 1C). One possibility is a high-dimensional representation in which the neural activity patterns for the four stimuli create an approximately tetrahedral geometry within the activity space. In this arrangement, all pairs of stimuli are separated by roughly equal distances. This geometry is unstructured and, hence, does not reflect any of the underlying structure or regularities of the stimuli. This geometry should contrasted to a low-dimensional representation, in which the four activity patterns are at the vertices of a square. Now the two patterns on each edge (e.g. the red and green fox) are closer than the patterns across the diagonal, which differ by two features. In this case, the structure of the world is reflected by the set of distances in the neural activity space.

To quantify these differences in representation geometry, two measures have been proposed: the parallelism score (PS) and the cross-condition generalization performance (CCGP) [1, 2]. Intuitively, both PS and CCGP quantify how consistent (or invariant) the representational direction for one semantic feature is across changes in the other features. Values of PS or CCGP close to 1 indicate that each semantic feature is represented in a dedicated direction, along which the representations are invariant to other features. In this subspace, the representation of a specific semantic feature are disassociated from the specific instances defined by different values of the other features, and hence are considered abstract. These representations are relatively low-dimensional (Fig. 1C). By contrast, in the high-dimensional (tetrahedral) geometry, PS and CCGP are close to 0. The PS and CCGP of a representation are fully determined by the representation kernel matrix, which quantifies the similarity between the neural activity patterns evoked by each pair of stimuli.

Both high- and low-dimensional representational geometries have been observed across brain regions. Abstract representations of task-relevant variables have been reported in multiple cognitive regions [1–7], and their emergence has been linked to the animals’ task performance [2].

On the other hand, many experiments show that the hippocampus (HPC) forms high-dimensional, sparse representations through pattern separation [8] and orthogonalization [9], which support memory discrimination and reduce interference. The HPC is strongly interconnected with the cognitive cortical regions mentioned above [10, 11]. How and why do these heterogeneous representational geometries emerge? Moreover, although it may seem intuitive that a high-dimensional, non-abstract hippocampal representation would hinder generalization, studies of patients with hippocampal or broader medial-temporal damage or dysfunction learning new semantic knowledge suggest a different picture [12–14]. These patients can acquire some new information, but their learning is typically much slower, and the resulting knowledge is often rigid and inflexible, with impaired generalization and transfer beyond the trained situations compared with controls. This deficit in generalization cannot be explained solely by a failure to form long term memory, because the patients do acquire the trained associations. These findings raise a central puzzle: why does disrupting the high-dimensional non-abstract hippocampal representation potentially impair the generalization behaviors, which likely depends on formation and use of low-dimensional abstract representations in other cognitive areas?

Here we investigate the question on how HPC may facilitate the formation of low-dimensional abstract representations and thereby support generalization, by using multi-regional circuit models trained on semantic-structured tasks. To isolate the contribution of hippocampus, we compare the representational geometry that emerges after training in a model with an explicit HPC module and in a matched model without it. We model HPC as a sparse, orthogonalizing autoencoder that forms high-dimensional representations of its inputs [15, 16].

We find that the impact of HPC on representation learning depends strongly on task structure, i.e. the representation geometry of the inputs and semantic knowledge of the task. When the input geometry is already aligned with the semantic space, removing the HPC module has relatively little effect. In contrast, when the input and semantic geometries are misaligned and the task requires a more complex input–output transformation, the HPC module plays a critical role in promoting abstract semantic representations in downstream areas and improving generalization. We interpret these misaligned tasks as those tasks requiring new schema formation, whereas aligned tasks are more schema-consistent and therefore depend less on HPC. Finally, we show that this HPC-dependent facilitation can be understood mathematically in terms of how the HPC module reshapes the loss landscape of the multiregional network during training. We demonstrate these findings across several settings. First, we study synthetic tasks in which input to semantic alignment can be varied continuously, and use feedforward network models trained on these tasks. We then show that the same principles extend to tasks with more naturalistic input statistics and recurrent network architectures. We further show that acquiring abstract semantic knowledge supports downstream, sample-efficient learning of flexible semantic-to-action mappings, especially under the systems consolidation framework which combines multiple pathways.

Together, our results describe circuit-level mechanisms by which HPC can support the emergence of low-dimensional abstract semantic representations and promote generalization across different stages of learning and different task scenarios. More broadly, our work provides an initial step toward understanding how multiple brain regions, equipped with distinct learning mechanisms, coordinate to achieve effective representation, yielding benefits that neither system learning system would have alone.

## Results

### A family of semantically-structured tasks that probe abstract representations

To probe how the hippocampus shapes abstract representations, we adopt a task-optimized network modeling approach commonly used in neuroscience [17–21] and investigate representational geometry in multi-regional neural network models trained on tasks in which both inputs and targets are generated from a latent semantic space (Figure 1B). Networks trained on similar tasks have been suggested to develop abstract representations of semantic variables in various scenarios [22, 23].

In this task, each stimulus input is associated with a set of binary semantic features, such as “animal/plant” and “real/imaginary” in Fig. 1. The desired output depends directly by these features. In the simplest case, the target output can be just the multidimensional binary semantic feature vector itself. More generally, we only require that the output geometry be aligned with the latent semantic geometry, as quantified by the representation kernel matrix (Fig. 1C, Methods). We show mathematically that whenever this alignment condition holds, the hidden representation learned after training is the same as in the case where the output vectors exactly match the semantic feature vectors. Consequently, the learned hidden representation does not depend on the specific way in which the output labels are generated from the semantic features (Methods, [22]).

This task setup admits two mathematically equivalent interpretations: (i) a single task with a multidimensional readout, such as reporting multiple stimulus attributes including color and shape; or (ii) a collection of related tasks performed over time, in which each task probes a subset of features, for example, reporting one feature of each stimulus across a sequence of tasks. Although these interpretations are equivalent and lead to the same training procedure, the latter is often used to model multitask learning in neuroscience [24,25] and machine learning [26,27]. This task family encompasses many binary label task paradigms studied in neuroscience, including both experimentally well-controlled settings [1–4, 28, 29] and more naturalistic, goal-directed tasks [30, 31]. Because we later consider multiple instances of this family, we refer to the overall setup as a “single” task throughout, and use “readout dimensions” to denote the different output components, whether they are read out simultaneously or across a sequence of related tasks.

Each task in this family is characterized by specific input and output geometries, defined by the representational kernel matrices in the input and output spaces. Although the output geometry is constrained to align with the latent semantic space, the input geometry can vary freely across tasks (Fig. 1D). For example, in one task, the binary semantic features might correspond to an object’s color and shape, while the inputs are pixel level images such as RGB values. In this case, stimuli that are closer in semantic space also tend to be closer in the input space: objects that share more semantic features also appear more visually similar. The input and output geometries are therefore aligned in this task (Fig. 2A, left). In another task, the inputs could be images of broccoli and foxes rendered in different colors with the binary semantic features to be animal/plant and real/imaginary as before (Fig. 2A, right). Unlike the color and shape task, semantic similarity in this setting does not imply visual similarity. Stimuli that are close in semantic space can appear very different in the input space.

**Figure 2:**
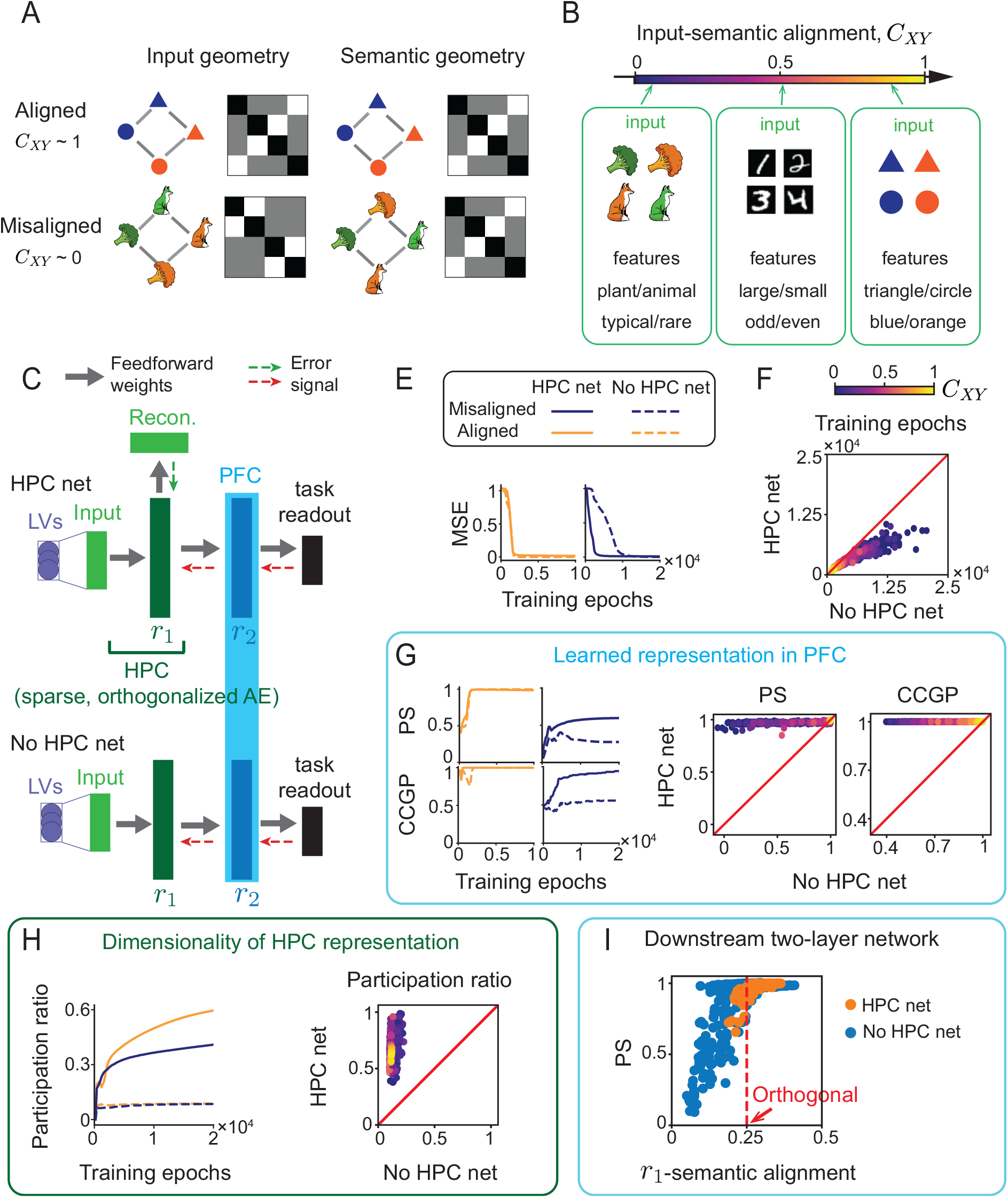
HPC facilitates learning abstract representations in multiregional networks. (A) Example aligned and misaligned input and output geometries, shown by stimulus configurations and their representation kernels. (B) Example tasks spanning the full range of input-semantic alignment *C*_*XY*_. Left: a misaligned task in which input similarity does not track semantic similarity. Middle: an intermediate-alignment task. Right: an aligned task in which input and output geometries are similar. (C) Architectures of the HPC net and the matched No HPC net. The HPC net contains an autoencoder-like HPC module with reconstruction loss and orthogonality regularization; the No HPC net replaces it with a purely supervised layer. (D) Example learning curves for aligned and misaligned tasks. (E) Learning time across tasks, comparing HPC net and No HPC net; points are colored by *C*_*XY*_. (F) Abstractness of learned PFC representations. Left, PS and CCGP during training for representative aligned and misaligned tasks. Right, PS and CCGP across tasks, comparing the HPC net with the No HPC net. (G) Dimensionality of the learned HPC/*r*_1_ representation, measured by participation ratio. Left, evolution during training. Right, comparison across tasks. The HPC net learns faster, forms higher-dimensional intermediate representations, and produces more abstract PFC representations, especially for low-alignment tasks. (H) In the downstream two-layer network, the alignment between *r*_1_ and semantic features is positively correlated with PS in the PFC layer.

Although these examples use visual images as inputs, the same task setup applies equally well to multimodal or sensorimotor inputs, and even to more abstract cognitive inputs, as long as they are associated with binary semantic features. Moreover, although we use static inputs here for simplicity, in practice, these inputs may also correspond to population activity in upstream cortical or subcortical areas. In that case, they can be viewed as time-integrated transformations of sensory and motor-related signals, and may also carry information about temporal history.

To better characterize differences across tasks, we quantify how closely stimulus representations in the input space align with those in the semantic or output space using the input-semantic alignment *C*_*XY*_. Specifically, *C*_*XY*_ is defined as the normalized inner product between the input and semantic representation kernels (Methods), and therefore takes values between 0 and 1. Because the output geometry is constrained to align with the latent semantic geometry, this input-semantic alignment also measures the alignment between input and output geometry. Intuitively, a smaller *C*_*XY*_ indicates a greater mismatch between input and semantic/output geometry and thus requires a more complex input–output transformation, whereas a larger *C*_*XY*_ indicates closer alignment and a simpler input-output mapping. To probe network models across a broad range of task structures, we generate a distribution of tasks with varying *C*_*XY*_ by systematically changing the input geometry (Fig.2B), using methods similar to previous work [23]. The resulting task distribution spans the full range of *C*_*XY*_ values (Fig. S1A) and provides a natural testbed for assessing how the hippocampus module affects learning across tasks with different levels of alignment or different levels of difficulty, as we will show below.

### HPC facilitates formation of abstract representations in the multiregional circuit

We now introduce the multi-regional network models used to train on these tasks and analyze representational geometry. To study the role of the hippocampus (HPC) in shaping representation geometry, we compare a network that includes an explicit hippocampal module, which we refer to as the HPC net, with an otherwise matched network that omits this module, which we refer to as the No HPC net (Fig. 2C).

In the HPC net, the hippocampal module is implemented as an autoencoder augmented with a regularization term that encourages orthogonality (decorrelation) of its neural activity patterns. In this framework, the reconstruction term in the autoencoder objective models the memory-related function of the HPC [32]. And the autoencoder architecture itself can be viewed as a neural network implementation of hidden state inference [33–35]. The orthogonality regularization is motivated by decorrelated codes reported across multiple hippocampal subregions, including dentate gyrus and CA3 [36, 37] and CA1 [9], and is also consistent with classical ideas [38] that the hippocampus performs an “expansion recoding” of multimodal sensorimotor inputs. We further find that this same orthogonality constraint promotes sparse activity in the HPC representation after task training (Fig. S1B). Related autoencoder models with related sparsity- or orthogonality-promoting regularization have been shown to produce place-field-like responses [15, 16] and autocorrelation statistics consistent with experimental observations [39]. Biologically, such orthogonality regularization in the hippocampus could arise from a combination of strong lateral inhibition in local circuits and behavioral timescale synaptic plasticity [40].

Downstream from the HPC module is a PFC module that receives inputs directly from the hidden layer of the HPC module and is trained in a supervised manner to produce the desired outputs (Fig. 2C). Similar supervised training has been widely used to model PFC representations in multitask settings [24,25,41]. Such supervised objectives may be implemented biologically through dopamine-dependent and other error or reinforcement signals. In contrast, the No HPC net has the same number of layers but differs in two key ways: (i) all layers are trained end-to-end using the supervised objective; and (ii) the first layer lacks orthogonality regularization and therefore does not develop sparse activity. (different legends for task readout/recons) These distinctions are reflected directly in the different terms of the training objectives for the two networks (Methods). Finally, the learning rate in the hippocampal layers of the HPC net, as well as the corresponding layer in the No HPC net (input → *r*_1_), is set higher than that of the remaining layers, reflecting the fact that the hippocampus is generally more plastic than the cortical areas.

We train both the HPC net and the No HPC net on this family of tasks with varying levels of input-semantic alignment. We assume that both networks operate in an overparameterized regime, where the number of neurons in each layer is large or comparable to the total number of stimuli in the task. After both networks achieve near-perfect performance, we analyze the geometry of neural representations across layers (Fig. 2C). All results shown below are obtained using the optimal regularization strength (Methods). However, we find that the qualitative conclusions are robust across a broad range of nonzero regularization strengths, provided that the regularization is not so large as to prevent the HPC net from learning the task (Fig. S1DE).

We first observed that the training dynamics of the two networks exhibit distinct signatures and depend strongly on the level of input-semantic alignment *C*_*XY*_ : for aligned tasks with large *C*_*XY*_ (simple task), both networks learn quickly (Fig. 2E, left). By contrast, for misaligned tasks with small *C*_*XY*_ (complex task), the HPC net learns substantially faster than the No HPC net (Fig. 2E, right). To characterize this advantage of HPC more systematically, we computed the learning times of each network, defined as the total number of training epochs required to reduce the mean squared error (MSE) on the training data to below a given threshold (Methods). Across tasks with different levels of difficulty (different values of *C*_*XY*_, Fig. 2F), we find that both networks learn more slowly as input-semantic alignment in the task decreases. However, the HPC net consistently shows a shorter learning time than the No HPC net, indicating more efficient learning (Fig. S1C). We note that this advantage is not simply due to the larger learning rate in the hippocampal layers, because the corresponding layer in the No HPC net is trained with the same learning rate. Interestingly, the training curves of the HPC net exhibit shorter plateaus, and in some cases, no clear plateau at all (Fig. S1G and Fig. 2E). Such plateaus are commonly associated with saddle points in the loss landscape that slow learning. These results, therefore, suggest that HPC improves learning efficiency by reducing or eliminating slow saddle points in the network’s loss landscape.

We next asked whether the more efficient training in the HPC net also leads to a more abstract representation of semantic features in the PFC layer. To test this, we computed the parallelism score (PS) and cross-condition generalization performance (CCGP), two standard measures on the abstractness of representation used in previous work [1, 2, 4, 28]. During training, both quantities increase in both network models as network performance improves (Fig. 2G, left). For aligned tasks, both networks achieve near-optimal PS and CCGP. In contrast, for misaligned tasks, the PFC layer in the HPC net exhibits substantially higher PS and CCGP than the No HPC net, indicating a more abstract representation of semantic features. This advantage of HPC is even clearer when PS and CCGP are plotted across tasks with different values of input semantic alignment *C*_*XY*_ (Fig. 2G, right). In the No HPC net, the PFC representation becomes progressively less abstract as input semantic alignment decreases and the task becomes more difficult. By contrast, in the HPC net, abstract representations consistently emerge across the full range of task difficulty, with both PS and CCGP remaining close to 1 even for tasks with low input-semantic alignment. Note that all the networks have achieved near-perfect performance but their PS and CCGP differ substantially.

To understand the reason for the robust emergence of abstract representation in the HPC net, we note that the main architectural difference between the two networks lies in the hippocampal layers. We therefore examined the dimensionality of neural representations in this layer in both networks, measured by the normalized participation ratio on the training data (Methods, Fig. 2H, left). The normalized participation ratio is defined as the participation ratio divided by the maximal dimension of the representation, a number ranging from 0 to 1. We find that although the No HPC net also increases its representation dimensionality during training, this increase is much smaller than that observed in the HPC net, for both aligned and misaligned tasks. Moreover, the dimensionality of the HPC net tends to be higher for tasks with low input-semantic alignment; the corresponding dimensionality in the No HPC net is relatively insensitive to *C*_*XY*_.

These trends become even clearer when the participation ratio is plotted across tasks with different levels of input-semantic alignment (Fig. 2H, right). Across the full range of tasks, the HPC net consistently forms a higher-dimensional representation of the stimuli in its HPC layer (*r*_1_ layer). Furthermore, this increase in dimensionality appears to be especially important for supporting abstract representations in misaligned tasks rather than aligned ones: we observe an increase in PS is correlated with an increase only in misaligned tasks (Fig. S1F). This observation is also consistent with the recent experimental data that schema-consistent knowledge can be acquired rapidly even in the absence of the hippocampus [42]. Together, these results suggest that the sparse, orthogonal representations formed in HPC may play a key role in enabling the emergence of low-dimensional abstract representations in downstream PFC, especially for tasks that require more complex input-to-output transformations, that is, tasks with low input-semantic alignment.

### Analysis of downstream two-layer network reveals the benefits of high-dimensional hippocampal expansion

Our simulation results suggest that high-dimensional hippocampal representations may play an important role in enabling the PFC to learn low-dimensional, abstract representations. At first glance, this may seem counterintuitive: if the goal is to form a low-dimensional abstract representation, one might expect it to be more helpful to preserve low dimensionality rather than first expand the representation into a higher-dimensional space. To better understand this phenomenon, we carried out a detailed mathematical analysis of the loss functions for both networks. We first note that, for a fixed representation geometry in the HPC layer (*r*_1_ layer), the abstractness of the PFC representation after training depends only on the representation kernel in the *r*_1_ layer and the representation kernel in the output layer (Methods). Motivated by this observation, we isolate the two-layer network downstream of the *r*_1_ layer, treating the *r*_1_ layer as the input layer in this reduced network.

We then examined how the abstractness of the learned PFC representation depends on the alignment between the representation kernels of the *r*_1_ layer and that of the output layer, which we refer to as the *r*_1_-*y* alignment. Plotting this quantity against the PS of the PFC representation after training (Fig. 2I), we find a threshold-like effect in the HPC net and the No-HPC net. When the *r*_1_-layer representation lies above this threshold, the learned PFC representation is nearly abstract. By contrast, when the *r*_1_-layer representation is below this threshold, the resulting PFC representation exhibits a broad range of PS values and is therefore not necessarily abstract. This threshold corresponds roughly to the alignment produced by a fully orthogonalized *r*_1_-representation. This observed threshold effect can be explained by directly computing the optimal solutions of the two-layer networks (Methods). Exploiting the convexity of the loss function of the reduced two-layer network, we show that whenever the *r*_1_-layer representation is more closely aligned to the output geometry than the fully orthogonalized representation,every global minimum yields the same PFC representation kernel, whose PS is exactly equal to 1.

These analyses help explain why the two network models produce different PFC representational geometries across tasks with different levels of input-semantic alignment *C*_*XY*_. When *C*_*XY*_ is high, the input geometry is already similar to the output geometry, so the task requires only a relatively simple transformation. In this regime, the network can simply preserve the input geometry across layers, so that the *r*_1_-layer, as well as the PFC layer, remain well aligned with the output and therefore to form abstract representations. As a result, in both the No-HPC net and the HPC net, the *r*_1_ representation typically has sufficiently high alignment with the output geometry to lie above the threshold described above, leading to abstract representations in PFC. In this case, orthogonalization in the hippocampal layer has relatively little effect.

By contrast, when *C*_*XY*_ is low, the task requires a more complex nonlinear transformation from input to output. In this regime, even though a purely supervised training (as in No HPC net) can still perform the task, it often fails to transform the misaligned input geometry into an *r*_1_ representation that is sufficiently aligned with the output. As a consequence, the *r*_1_-*y* alignment may remain below the threshold, and the downstream PFC layer does not reliably develop an abstract representation. In the HPC net, however, the orthogonal dimensional expansion in the hippocampal layer (in the HPC net) effectively “erases” those misaligned low-dimensional structures in the input and pushes the *r*_1_ representation to the threshold, thereby enabling the emergence of an abstract representation in PFC after training.

The less obvious part of this explanation is that a fully orthogonal representation, which might intuitively seem unstructured and therefore not aligned to any low-dimensional structure, turns out to lie just above the *r*_1_-*y* alignment threshold required for abstract representations to emerge. This seemingly counterintuitive result can be understood more intuitively by examining the geometry of the space of representation kernels, which are positive-definite matrices. In Euclidean space, a random unstructured vector in high dimensions is approximately orthogonal to any fixed vector. By contrast, in the space of positive semidefinite matrices, a representation kernel associated with an unstructured geometry has a strictly positive inner product with any other nonzero representation kernel. In this sense, a fully orthogonalized representation is not aligned with any particular low-dimensional structure in the original Euclidean space, but remains positively aligned with any low-dimensional geometries in kernel space, including the one defined by the semantic features in our task. Since the optimal hidden representation learned by the downstream network depends on the input and output kernels rather than on the original Euclidean coordinates of the input and output representations (Methods), this positive alignment in kernel space facilitates the formation of low-dimensional abstract representations downstream. Thus, the fully orthogonalized kernel occupies a special position in the space of representation kernels: it suppresses misleading low-dimensional structure in the input while retaining positive alignment with the semantic structures in the output. This special role explains why the sparse, high-dimensional expansion performed by the hippocampus can facilitate, rather than hinder, the learning of low-dimensional abstract representations in downstream circuits.

### HPC facilitates learning of abstract representation in naturalistic data and in recurrent networks

We have previously used synthetic tasks to probe the role of HPC, with within class variability for each binary latent class generated from Gaussian distributions. This controlled setup allows us to systematically vary input-semantic alignment and examine its effects. However, realistic data often exhibit within class variability that is not well captured by Gaussian distributions. In addition, the network models studied above assume that computation in PFC, or more generally in semantic cortical areas, can be approximated by feedforward networks, which allows us to analytically characterize the global optimum of the downstream two layer loss. Nevertheless, we find that the advantages of HPC identified in this simplified setting persist even when these assumptions are relaxed, both when within class variability departs from Gaussian structure and when the PFC module performs recurrent computations across time.

To test whether high dimensional HPC representations provide similar benefits under realistic within class variability, we construct analogous semantically structured tasks using the Dsprites and 3DShapes datasets (Fig. 3A-C) and evaluate the role of HPC by comparing the two network models. To illustrate how we construct tasks from these dataset to probe the abstract representation, we take the 3Dshapes dataset as an example. In these image dataset, each image is assigned to multiple continuous latent variables related to visual attributes, including shape, size, orientation or colors of the object and background in the image (Fig. 3B). To adapt this dataset to the class of tasks considered previously (Fig. 1-2), we first binarize the continuous visual latent variables to obtain binary visual latent features (*l*_1_, *l*_2_, .., *l*_*D*_ in Fig. 3B). As in our earlier task formulation, we then define binary semantic features for each image so that they depend only on these binarized visual latent features. These visual to semantic mappings are generally conjunctive with respect to the latent visual features (Methods). Examples of such semantic features include whether an image has high or low contrast, which depends on both object and background colors, and whether it contains a warm cylinder or a cool cube, which depends on both object shape and color (Fig. 3B). As before, the multidimensional task readout **y** for each image is assumed to be aligned with the semantic features, while the task input to the network is the pixel level representation of the image. By varying the visual to semantic mapping, we generate a family of tasks with different levels of input-semantic alignment *C*_*XY*_, corresponding to semantic features that depend on different aspects of the visual latent variables (Methods). For each task and its corresponding visual to semantic mapping, we randomly sample the same number of training examples from each binary semantic class, so that the overall output geometry remains aligned with the abstract geometry of the semantic space.

**Figure 3:**
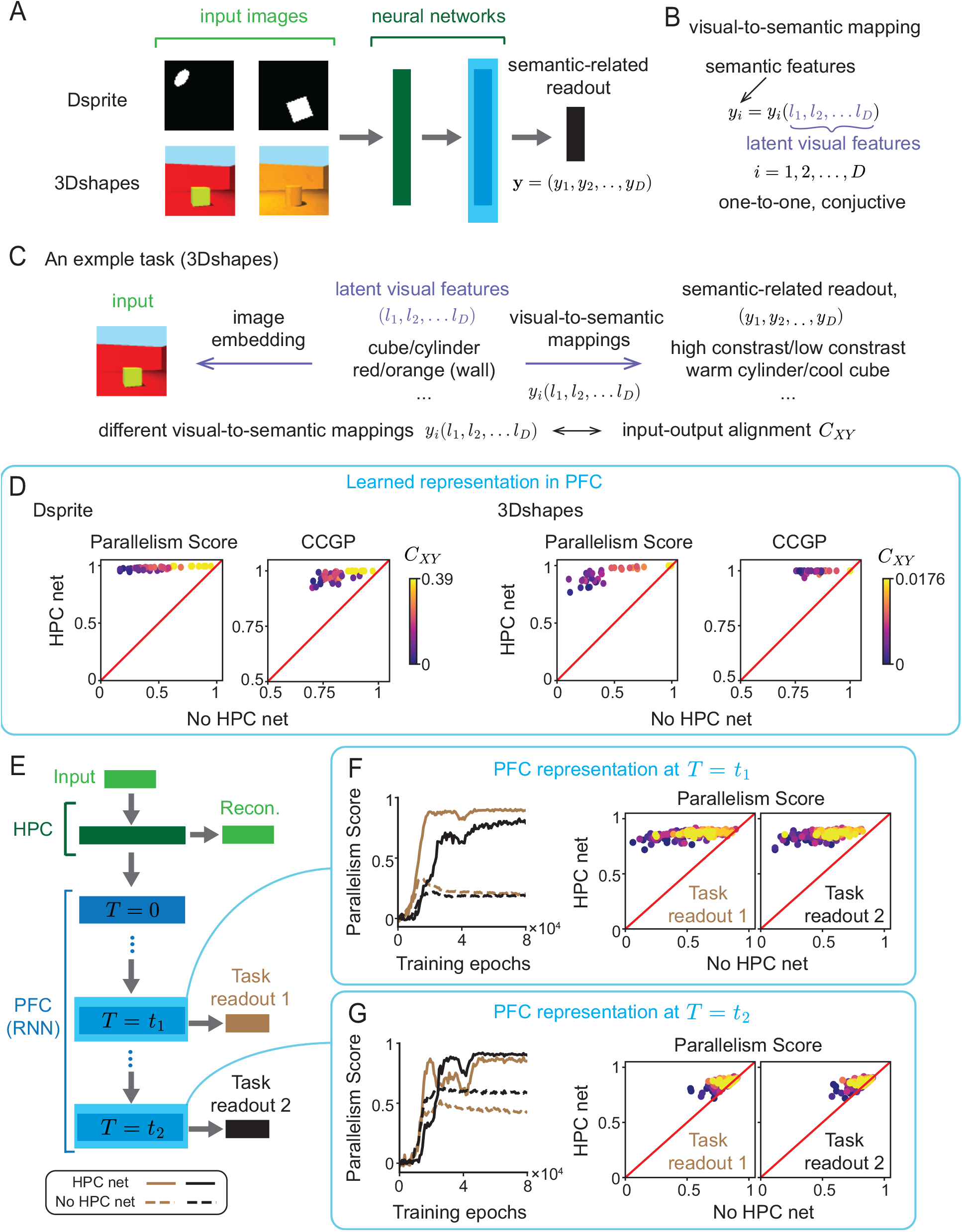
Advantages of HPC module extend to naturalistic inputs and recurrent downstream computation. (B) Construction of tasks from the Dsprites and 3DShapes datasets. Images are generated from latent visual features, which are binarized and mapped conjunctively to semantic features that define the task readouts. (C) Example task from 3DShapes, illustrating how different visual-to-semantic mappings induce different levels of input-semantic alignment *C*_*XY*_. (D) Abstractness of PFC representations learned from naturalistic image inputs. For Dsprites and 3DShapes, PS and CCGP are shown for the HPC net versus the No-HPC net across tasks, colored by *C*_*XY*_. (E) Recurrent-network version of the model, in which the PFC module is an RNN and different semantic readouts are required at times *t*_1_ and *t*_2_. (F,G) PS for semantic features during training for PFC representations at *t*_1_ and *t*_2_. In both naturalistic and recurrent settings, the HPC net continues to promote more abstract downstream representations, with the strongest advantage for low-alignment tasks.

We train both the HPC net and the No-HPC net on such tasks generated from both Dsprites and 3Dshape dataset. After each network reaches near perfect performance, we evaluate the abstractness of the PFC representation by computing PS and CCGP with respect to the semantic features (Fig. 3D). Similar to the synthetic tasks studied earlier (Fig. 2), when *C*_*XY*_ is high, the PFC representations in both the HPC net and the No-HPC net exhibit high PS and CCGP, indicating abstract representations of the semantic features. As *C*_*XY*_ decreases, however, PS and CCGP of the PFC representation in the No-HPC net also decrease, whereas both quantities remain close to 1 in HPC net. We nevertheless observe two subtle differences relative to the synthetic tasks (Fig.2G, right). First, the abstractness of the PFC representation in the HPC net shows a slight decline as input semantic alignment decreases. More specifically, the two abstractness metrics show different sensitivities across the two image datasets: CCGP is more sensitive in Dsprites, whereas PS is more sensitive in 3DShapes. These differences may reflect differences in within class variability associated with the underlying 2D and 3D spatial structure of the datasets. Second, because of the preexisting relationship between pixel level inputs and the latent visual features, the task generation process used here does not span the full range of possible input semantic alignment values. Despite these differences, the results consistently show that the HPC net learns more abstract semantic representations than the No HPC net across all levels of input semantic alignment achieved in these datasets.

Many previous observations of the abstract representations of task-relevant variables have come from the cognitive cortical areas, such as the prefrontal cortex [1–3], the orbitofrontal cortex and the anterior cingulate cortex [1, 43]. Some of these regions are known to contain strong recurrent excitatory connections and to perform nontrivial recurrent computations, such as working memory, that process input information over time. To investigate how such recurrent processing influences the results obtained with the feedforward PFC module above (Fig. 2), we replaced the PFC module with a recurrent neural network that receives input from the HPC module in the HPC net (Fig. 3E) and is trained in a supervised manner. To study how the network maintains and processes information over time, we also modified the synthetic tasks used above so that different components of the semantic features are read out at two separate time points within each trial, while the input is presented only at the beginning of the trial. The No HPC net is modified in the same way, except that all layers are trained in a supervised manner.

We train both networks on these modified tasks with different levels of input-semantic alignment *C*_*XY*_. After training, we assess the abstractness of the PFC representation by computing the PS at the two output time points with respect to both sets of semantic features, including those read out at that time point and those read out at the other time point. For misaligned tasks, the HPC net again develops a more abstract representation than the No HPC net at both readout times (Fig. 3FG), consistent with the feedforward case. Moreover, in both networks, the semantic features read out at a given time point are represented more abstractly than those read out at the other time point. Across the full range of input semantic alignment values, the PS of the PFC representation remains consistently higher in the HPC net than in the No HPC net. Unlike in the feedforward setting, where the HPC net often learns an almost perfectly abstract representation with PS close to 1 (Fig. 2), the gradient optimizer is less effective in the recurrent PFC network and therefore produces greater variability in PS, even in the HPC net. Nevertheless, these results show that the advantage of HPC in promoting downstream abstract representations persists across tasks that require temporal processing and recurrent computation.

### Abstract semantic representation builds in inductive bias to support flexible semantics-to-action mappings

HPC has been proposed to form high-dimensional, sparse representations of external stimuli; such representations are often expected to support fine-scale discrimination at the expense of generalization. Because generalization requires the system to ignore fine-scale details of individual stimuli, extract their shared structure and apply that shared structure in novel scenarios. Such a trade-off generally holds for a single-module network with homogeneous neural firingrate sparsity [44]. At the same time, previous experiments have reported the observation that compared with normal people, hippocampus lesion patients show slower learning of new semantic concepts and, even after learning, are impaired in applying those concepts in novel contexts (for example, when asked to use a newly learned word in a new sentence). The acquired concepts also tend to remain tied to the specific training context. Our results suggest a possible resolution to this puzzle. We find that the presence of an HPC module helps downstream semantic cortical areas form abstract representations of semantic features more effectively. Because this type of low-dimensional representation has been proposed to support out-of-distribution generalization [22, 28], our findings point to a potential mechanism by which HPC may promote generalization in multi-module networks.

To illustrate how the HPC net supports such generalization in a novel scenario, we consider a two-phase training paradigm. In Phase 1, both networks, the HPC net and No HPC net, are trained on the same family of tasks with binary semantic features described above (Fig. 3A, left). We focus specifically on misaligned tasks with low input–output alignment. After Phase 1 training, we expect the PFC layer in the HPC net to develop an abstract semantic representation of the stimuli, whereas the No HPC net does not. These learned abstract representations can be interpreted as a structured scaffold of semantic knowledge, or a schema [45, 45, 46], that can guide behavior in future tasks. To test whether such a schema facilitates generalization, we introduce Phase 2 training. In this phase, we take the learned stimulus representations in PFC as input and train a new motor output pathway to solve a novel task (Fig. 3A, right). Crucially, the outputs in the new task are fully determined by the semantic features acquired during Phase 1. We consider two broad classes of such transfer tasks below.

The first class of such tasks consists of those in which the motor output depends only on a subset of the semantic features and is invariant to the remaining ones. To produce the correct action, the motor pathway must therefore learn to ignore the irrelevant semantic features. Two examples in this class are the majority and parity tasks (Fig. 3B). In the majority task, the network output is formally defined as the sign of the sum of the binary features (Fig. 3B). Intuitively, the network must report the label corresponding to the majority of the relevant semantic features. For example, consider a task in which each input is an image containing three objects, each of which can be either a circle or a triangle, possibly with different sizes and orientations that are irrelevant features to the task. The correct output is then the shape represented by the largest number of objects. In the parity task, the network output is formally defined as the product of the binary features (Fig. 3B). Here, the motor output depends on whether an odd number of binary features are present. One example is again a task in which each input contains three objects, each either a circle or a triangle. The correct output is the shape that does not form a pair. We note that, in the semantic space, the majority task is linearly separable, whereas the parity task is not.

To test generalization in these two tasks, we train the motor pathways using randomly sampled stimuli with fixed irrelevant features, and then evaluate performance on novel stimuli in which the irrelevant features are changed (Fig. 3B). In the three object example, this corresponds to training on blue circles and triangles and testing on red ones. This therefore constitutes a form of out of distribution generalization. After Phase 2 training, we find that the learned motor pathways in both networks generalize better as the number of training samples increases. However, the HPC net consistently outperforms the No HPC net across all training set sizes, with the advantage being especially pronounced for the nonlinear parity task (Fig. 3C). Moreover, even when trained on all stimuli in the training set, the No HPC net still shows a fundamental difficulty in ignoring the irrelevant features. Based on the differences between the two networks observed above, we hypothesized that this superior performance of the HPC net arises from its effective formation of abstract representations during Phase 1 training. Consistent with this idea, we observe positive correlations between the abstractness of the PFC representation and out of distribution generalization performance in both networks for both tasks (Fig. 3C). Analytical calculations in simple two-layer linear networks further indicate that the ability to ignore irrelevant features is rooted in representing relevant and irrelevant semantic variables in orthogonal subspaces (Methods).

Similar results are observed for a second class of tasks that requires the network to generalize compositionally [47–49], rather than simply ignore irrelevant features. In these tasks, the new motor output is generated by partitioning the semantic features of each stimulus into two groups: stimulus features and rule features (Fig. 4D). The two examples we consider here are the context-dependent majority-minority task and the context-dependent reversal task, each with a single rule feature. In the context-dependent majority-minority task, the motor output is the same as in the majority task when the rule feature is +1, and is flipped when the rule feature is *−*1. In the context-dependent reversal task, the motor output either copies the stimulus features or reverses them, depending on the value of the rule feature. In the image example with three objects, both tasks can be interpreted as adding a fourth object whose shape serves as a contextual cue that determines how the motor output should be generated from the features of the original three objects.

**Figure 4:**
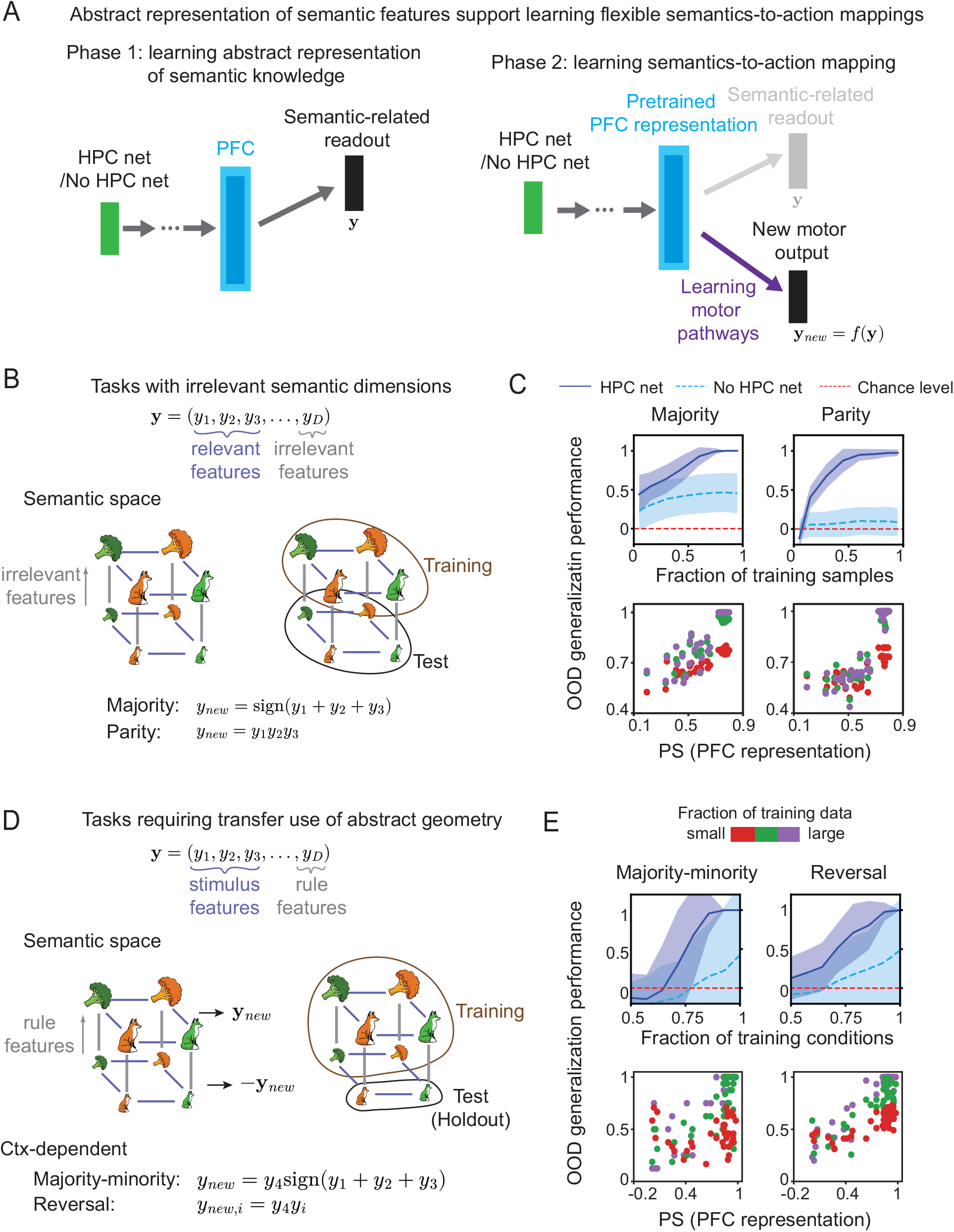
Abstract semantic representations support flexible semantics-to-action mappings. (A) Two-phase training paradigm. In phase 1, the network learns semantic-related readouts and forms a PFC representation of semantic structure. In phase 2, a new motor output pathway is trained on top of the pretrained PFC representation to solve a novel task. (B) Transfer tasks with irrelevant semantic dimensions. (Left) Examples of majority and parity tasks, in which the correct motor output depends on relevant semantic features and should generalize across changes in irrelevant features. (Right) Training/Test split used to assess out-of-distribution generalization. (C) Generalization performance versus fraction of training samples, and its correlation with PS of the PFC representation. The Pearson correlation for different fraction of data are *r* = 0.47 (small), *r* = 0.68 (intermediate) and *r* = 0.66 (large). (D) Transfer tasks requiring compositional use of abstract geometry. (Left) Examples of context-dependent majority-minority and reversal tasks, in which a rule feature determines how stimulus features should be mapped to action. (Right) Training/Test split with hold-out conditions. (E) Generalization performance versus fraction of training conditions, and its correlation to PS of the PFC representation. Pretrained abstract PFC representations in the HPC net provide an inductive bias that enables more sample-efficient out-of-distribution generalization. The Pearson correlation for different fraction of data are *r* = 0.73 (small), *r* = 0.86 (intermediate) and *r* = 0.89 (large)

During Phase 2 training, we randomly select a subset of semantic classes and hold out all stimuli belonging to those classes, while training the motor pathways on the remaining stimuli. As a result, the training set does not fully cover the semantic space. After training, both networks are tested on the hold-out stimuli to determine whether they can generalize compositionally in these tasks (Fig. 4E). We observe a similar advantage of the HPC net in this second class of tasks: its generalization performance is substantially higher than that of the No HPC net. Moreover, when we plot the PS of the PFC representation against generalization performance, we find a strong positive correlation, provided that the number of training conditions is not too small.

These results show that the formation of abstract representations provides an inductive bias that enables downstream motor pathways to learn flexible semantic-related actions and support sample-efficient out of distribution generalization. Given the role of HPC in facilitating the effective learning of such abstract representations from the outset of learning (Phase 1), our findings may offer a possible explanation for why hippocampal lesion patients exhibit deficits in generalization behavior.

### Coordination of HPC-dependent and HPC-independent pathways supports input-based generalization

The results above show that the formation of abstract representations in PFC enables the network to achieve sample-efficient out-of-distribution generalization when trained on new tasks whose outputs depend on semantic features. This advantage arises because the network can reuse the learned representational geometry formed during Phase 1 for stimuli encountered there. Such reuse is effective when test inputs are similar to those seen during phase 1 training. However, when a test input differs substantially from all training inputs, the network may no longer be able to rely on the abstract scaffold formed in PFC during Phase 1. Moreover, because the high-dimensional expansion in the HPC module decorrelates the inputs, it destroys the low-dimensional structure in the input space and therefore becomes less able to generalize on the basis of input similarity, relative to the No HPC net. The brain is known to contain both types of pathways: pathways that go through HPC and pathways that do not, analogous to the HPC net and the No HPC net. Motivated by complementary learning systems theory [38, 50–54], we therefore consider a network that consists of both pathways and investigate whether combining them can improve input-based generalization (Fig. 5A).

**Figure 5:**
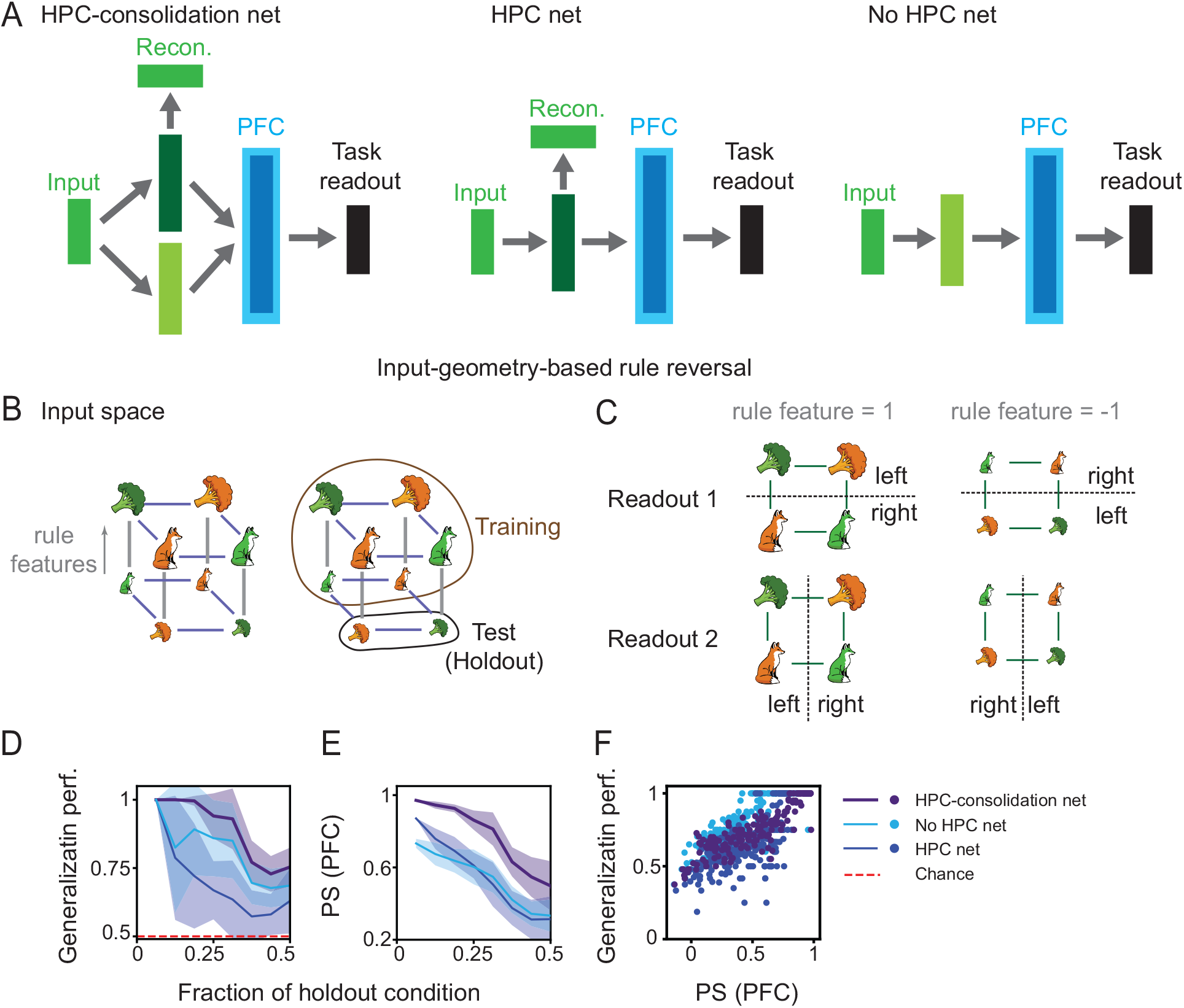
Combining HPC-dependent and HPC-independent pathways improves input-based generalization. (A) Architectures of the HPC-consolidation net, HPC net, and No HPC net. The HPC-consolidation net uses an additional HPC-independent pathway that is trained to match the PFC representation generated through the HPC-dependent pathway. (B) Input-geometry-based rule-reversal task. The plots show the input geometry and training/test split in input space with held-out conditions. (C) The two readouts are induced by the rule feature. This task probes generalization that depends on preserving input structure. (D-F) Comparison of generalization and representation geometry across models. The HPC-consolidation net achieves the best generalization performance and the highest PS among the three networks. (D) Out-of-distribution generalization performance versus fraction of held-out conditions. (E) PS of the PFC representation versus fraction of held-out conditions. (F) Positive correlations between generalization performance and PS in all three networks. Pearson correlations: *r* = 0.92 (HPC-consolidation net), *r* = 0.77 (No HPC net) and *r* = 0.77 (HPC net).

The most straightforward way to train the two-pathway network on a task would be to use end-to-end backpropagation from the output layer. However, because the learning rate in the HPC module is set much higher than that in the No HPC pathway, the network will end up relying almost exclusively on the HPC-dependent pathway to perform the task, while the HPC-independent pathway remains largely untrained. In this regime, the full model is equivalent to a single pathway network dominated by the HPC route. This phenomenon is known in the deep learning literature as neural race reduction [55]. The intuitive explanation is that, when multiple pathways are driven by the same behavioral error signal, the fastest pathway can rapidly learn to produce the correct output, after which the shared error becomes too small for the other pathways to learn effectively.

To overcome this problem, we introduce a memory consolidation mechanism for the HPC-independent pathway: instead of being trained solely on task outputs, this pathway is trained to match the PFC representation generated by the HPC-dependent pathway (Methods). We refer to the resulting model as the HPC consolidation net. Biologically, such a consolidation mechanism could be supported by offline replay previously observed in HPC and PFC [56]. Moreover, after training, the HPC-independent pathway alone is sufficient to generate the correct task output, so performance remains intact even after lesioning the HPC module. This eventual HPC independence of learned knowledge is consistent with behavioral findings from patients with classical hippocampal lesions [14, 57].

Because we are primarily interested in out-of-distribution generalization based on input structure, we focus on the context-dependent reversal task, in which the abstract semantic representation is embedded in the input space by flipping the semantic labels according to a rule feature (Methods). The task output is aligned with the original semantic features (Fig. 4B). Intuitively, when the rule feature is +1, the output aligns with the input, whereas when the rule feature is *−*1, the output is flipped. Because the original abstract semantic structure is partially preserved in the input space, we expect that networks trained on a randomly sampled subset of inputs can generalize to hold-out inputs with different semantic features by exploiting the low-dimensional structure in the input space.

We compare generalization performance across three networks: the HPC-consolidation net, the HPC net, and the No HPC net. After training, all three networks generalize above chance level (Fig. 5C, left and Fig. S2A). Among them, the HPC-consolidation net consistently achieves the best generalization performance across different numbers of training conditions. The No HPC net outperforms the HPC net, perhaps because it preserves more of the preexisting structure in the input space.

We next computed PS to quantify the abstractness of semantic representations in PFC across all stimulus conditions, including the holdout conditions (Fig. 5C, middle). For the HPC-consolidation net, the PFC representation was evaluated based on the activity generated via the HPC-independent pathway after consolidation. Consistent with their generalization performance, the HPC-consolidation net shows higher PS than both the No HPC net and the HPC net. More importantly, across all three networks, generalization performance is positively correlated with the abstractness of semantic representations in PFC after training.

These results suggest that the HPC-consolidation net can learn abstract semantic representations more effectively than either single pathway networks, by first constructing a partial representation scaffold through the HPC-dependent pathway and then consolidating it into the HPC-independent pathway. This more effective formation of abstract representations in turn supports better out-of-distribution generalization based on input structure. Notably, the advantage of combining the two pathways is specific to out-of-distribution generalization and is not observed for simpler in-distribution generalization (Fig. S2B).

## Discussion

Neural activity in many brain areas has been found to encode the same set of cognitive variables. Yet these variables are often represented with different geometries across regions, allowing the brain to flexibly support different behavioral demands, such as fine discrimination versus generalization [58]. In particular, the hippocampus and cognitive cortical regions have been suggested to exhibit high-dimensional orthogonal representations and low-dimensional abstract representations, respectively. Here, we investigate how these distinct representational geometries interact across brain regions, and specifically how high-dimensional hippocampal representations shape low-dimensional representations in PFC. More broadly, our work provides an initial step toward understanding how different representational geometries across brain regions interact during learning.

### High-dimensional hippocampal expansion as a route to abstraction

We find that, although the high-dimensional expansion performed by HPC appears to move in the opposite direction of forming a low-dimensional representation, it in fact removes the low-dimensional structure in the input that is misaligned with the semantic structure. This enables downstream areas to more effectively learn low-dimensional abstract representations from semantic-driven behavioral readouts. It may seem surprising that a high-dimensional orthogonal HPC representation, which is not itself aligned with any particular low-dimensional geometry, can nevertheless facilitate the formation of abstract representations. We show that this phenomenon can be understood mathematically from the special geometry of the space of representation kernel matrices and from the structure of the loss landscape of deep neural networks.

Although our mathematical analysis assumes a perfectly orthogonalized HPC representation, the HPC layer has a finite number of neurons and limited capacity, and therefore must perform some forms of memory compression. Such compression introduces correlations between the representations of different stimuli and can give rise to a specific low-dimensional structure. We indeed observe evidence of such structure in some simulations: in many misaligned tasks, the participation ratio of the HPC representation after training is substantially smaller than that of a fully orthogonal representation (Fig. 2), indicating that representations of different stimuli remain correlated. In the training objective of the HPC module, both the reconstruction term and the supervised learning signal at the HPC input layer may contribute to this compression effect and thereby to the formation of low-dimensional structure in HPC. Despite the presence of this residual low-dimensional structure, the representation in HPC remains consistently higher-dimensional than that in PFC after training, even for misaligned tasks. This pattern does not occur in the No-HPC net, where the *r*_1_ layer and the PFC layer have similar dimensionality. Together with the corresponding differences in PS and CCGP, these differences in dimensionality suggest that the overall effect of expansion in the HPC layer, even when it is not perfectly orthogonal, can still robustly promote the formation of low-dimensional, abstract semantic representations downstream.

While our work focuses on the high-dimensional expansion in the hippocampus, the computational mechanisms for the interactions between representation geometries revealed here may apply more broadly to other brain regions that perform related forms of expansion coding, such as primary sensory cortices [59] or the cerebellum [60]. It is therefore possible that high-dimensional expansion in these regions also contributes to the formation of low-dimensional abstract representations in downstream areas through mechanisms similar to those identified here. Compared with regions specialized primarily for sensory or motor processing, however, HPC may be unique in performing such expansion recoding for multisensory, sensorimotor, and cognitive inputs.

Compared to these other brain regions specialized in sensory or motor processing alone, HPC is probably special in performing such expansion recoding for multisensory or sensorimotor inputs and cognitive variables. This view is supported by experimental findings showing that hippocampal lesions impair discrimination between different contexts containing rich multisensory information, pointing to a distinctive role of HPC in processing this type of input. Because much semantic knowledge depends on multimodal inputs embedded in rich contextual structure, our results suggest that HPC may play an especially important and potentially irreplaceable role in the formation of generalizable semantic knowledge.

### Relations to other multiregional circuit model

Our results are closely related to complementary learning systems theory [50–52], but differ from it in important ways. Complementary learning systems theory proposes that HPC forms separated representations of individual experiences to support memory discrimination, whereas neocortex forms more overlapping representations that extract statistical regularities from the environment. To mitigate catastrophic interference in continual learning settings, these distinct representational forms are thought to be paired with fast learning in HPC and slower learning in cortex. Although the models studied here are consistent with this general picture, our focus is different. We ask how representational geometry in HPC shapes representational geometry in cortex in the multitask settings commonly used in neuroscience and machine learning [22, 24–26], rather than whether HPC reduces catastrophic interference during continual learning. In our setting, the benefit of HPC is to reshape the downstream loss landscape so that the learned downstream representation becomes aligned with the semantic structure of the stimuli. Indeed, in all our results, the No-HPC net can still achieve near perfect task performance and does not suffer from catastrophic interference. We further show that this benefit depends strongly on task structure: HPC is most helpful for difficult tasks with low input-semantic alignment, but is not equally important for all tasks. This effect of HPC on shaping representational geometry also extends to continual learning scenarios in which different tasks are learned sequentially (Fig. S3A). Finally, inspired by complementary learning systems theory, we introduce a consolidation mechanism to address the issue of input-based generalization. Notably, after consolidation, the HPC-consolidation network with two pathways generalizes better than either single pathway network alone. This advantage again arises from learning a more semantically-aligned representation geometry, rather than from avoiding catastrophic forgetting, since all three networks achieve near perfect training performance after learning (Fig. 5).

Another line of modeling work related to our study concerns interactions between entorhinal cortex and hippocampus, such as the Tolman-Eichenbaum Machine and vectorHASH [61, 62]. In these models, HPC supports generalization by binding novel sensory inputs to existing low dimensional scaffolds. These scaffolds are often associated with grid cell modules in entorhinal cortex and are largely hard-wired during development. By contrast, the generalization advantage of HPC in our work arises primarily from interactions between representational geometries across different brain regions, rather than from successful HPC binding. Moreover, our focus is on population-level neural geometry rather than on single-neuron tuning properties, which are often emphasized in these models. Although we do not explicitly test whether the same hippocampal binding mechanism also gives the HPC net an advantage over the No-HPC net, preliminary simulations in sequential learning settings suggest that sparse, high dimensional hippocampal representations may also support more effective binding based generalization (Fig. S3B).

### Experimental predictions

Our results provide a potential mechanism for explaining the deficits observed in generalization behavior when HPC lesion patients learn novel concepts [14]. These results also make novel predictions on representation geometries after HPC lesion, for these novel concept learning that shows generalization deficit, the neural representation of the related semantic cortical areas will be less abstract compared to controls. As more and more experiments on training animals to solve semantically meaningful and complex tasks come up in the future, we expect these predictions to be tested by recording from HPC lesion animals.

### Future direction

While we here focus on the interaction between representation geometries from the perspective HPC → PFC, HPC and PFC in the brain are bidirectionally connected. Low-dimensional representation geometry in PFC may also impact memory contents retrieved from the HPC as well as what new memory gets formed and added in the HPC [63, 64]. This bidirectional exchange of information between HPC and PFC may be crucial for online closed-loop behaviors such as contextual inference and planning in semantically-rich environments, which would be an important future direction. Our work thus represents a first step towards understanding how such interconnected brain areas coordinate to learn semantic knowledge by forming heterogeneous representation geometries and supporting flexible generalization across diverse novel situations.

## Methods

### Task models

#### Synthetic tasks for probe semantic representation

There are in total *P* training data points, 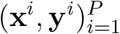. And each training data (or training stimulus) is associated *D* binary labels, 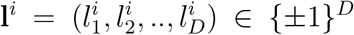, representing the *D* latent semantic features for the stimulus. Based on the semantic features, all stimuli are divided into 2^*D*^ classes, and each class has an equal number of training stimuli.

The output **y**^*i*^ for each stimulus only depends on its latent label **l**^*i*^, **y**^*i*^ = *f* (**l**^*i*^) (semantic-related readout). Moreover, we require the output kernel matrix of the training data to be aligned with the kernel matrix in the latent space,

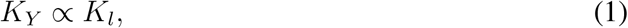

where the kernel matrices are defined as

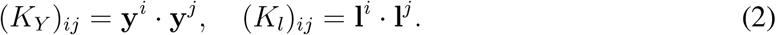

We note that this alignment condition does not fully constrain the choice of the semantic-output map *f*. However, because the hidden representation after training in both HPC net and no-HPC net (See section “Optimal representation only depends on the input-output kernel” below) only depends on the input and output kernel matrices, different choices of *f* satisfying Eq.(1) would all yield the same hidden representations.

There are a few natural choices for *f*. One is simply the identity map, **y**^*i*^ = **l**^*i*^ ∈ ℝ^*D*^, which also holds in our simulation. Another typical choice is to assume that behavior readouts are random linear readouts of the linear space [22]. In this case, the *k*th output dimension is assumed to be a random projection of the latent label, 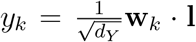, where the component of **w**^*k*^ is i.i.d. sampled from Gaussian distribution with mean zero and unit variance. *d*_*Y*_ is the total number of output dimensions. With a sufficiently large number of behavior readouts (each readout can be thought of as a different semantic-related task, as we mentioned in the main text), the output kernel matrix is

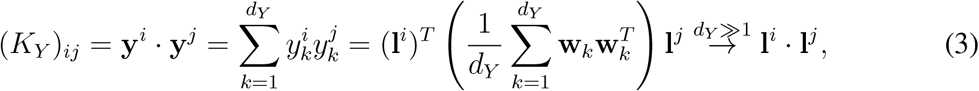

which aligns with the latent kernel matrix. For a general nonlinear readout of form 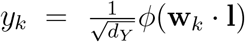 with smooth *ϕ* and *ϕ*(0) = 0, the output kernel matrix can be written as

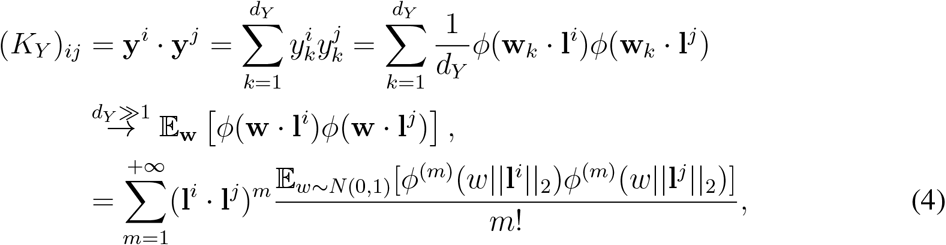

where the last line uses Gaussian-Hermite expansion and *ϕ*^(*m*)^ is the *m*th derivative of *ϕ*. If all its derivatives are bounded by a constant, the high-order terms in the summation decays quickly ~ (*k*!)^*−*1^ and the sum will be dominated by the 1st term, which is aligned with the latent kernel *K*_*l*_.

On the other hand, the input **x**^*i*^ for each data is generated to have a diverse range of input-semantic alignment. For a task, we first generate the input kernel matrix as follows

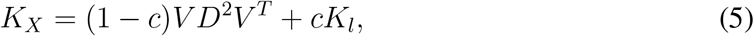

where *c* is sampled uniformly from [0, 1] and *V* ∈ ℝ^*P ×l*^ is a random matrix with orthonormal columns and *D* ∈ ℝ^*l×l*^ is a diagonal matrix whose diagonal vector is randomly sampled uniformly from a sphere. *l ≤ P*. We choose the radius of the sphere such that tr*D*^2^ = tr*K*_*l*_. Intuitively, the above input kernel is a random perturbed version of the kernel in the semantic space. The amount of random perturbation is controlled by *c*. For each randomly sampled *K*_*X*_, we compute its input-semantic alignment based on the following expression,

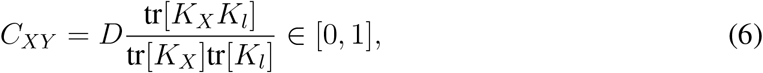

where the normalization factor *D*, which equals the number of latent features, is to ensure that *C*_*XY*_ = 1 when *K*_*X*_ = *K*_*l*_. Here we measure the alignment using uncentered kernels rather than centered ones as in many centered kernel alignment analysis. This is because the optimal representation learned by the network models depend on the uncentered kernels rather than the centered ones (section “optimal representation only depends on the input-output kernel”).

We divide the interval [0, 1] evenly into ten subintervals and sample many random *K*_*X*_ such that the number of tasks within each interval is 50. From *K*_*X*_, we generate input data with this input kernel. As we will show later (section “optimal representation only depends on the input-output kernel”), the optimal representation in the network depends only on the input and output kernel matrices, so it doesn’t depend on how we generate the input data. These are the families of tasks we tested the HPC net and the no-HPC net on in Figure 2.

#### Dsprites and 3Dshape generated task

For the Dsprites dataset, we take 4 latent visual features from each image, representing shape, scale, orientation and *X* position of the image. The remaining latent feature (the object’s *Y* position in the image) is fixed. Except for the shape feature, all the other latent visual features are continuous. For the shape feature, we choose images that are either square or elliptical. For the continuous features, we binarized them. In total, we obtain 4 binary labels (*l*_1_, *l*_2_, *l*_3_, *l*_4_) for each image. The semantic features (which are also the output labels for each image) are defined as a one-to-one function from 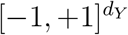 to itself,

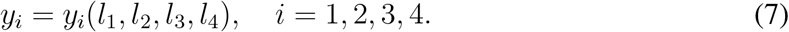

As this visual-to-semantic mapping is one-to-one, it can be represented as a permutation of the vertices of a *d*_*Y*_-dimensional hypercube. To generate a family of tasks with different visual-to-semantic mappings, we randomly sample such permutations, each of which corresponds to a task with the newly generated semantic features.

The network’s input is the image’s pixel representation (greyscale representation). The input-semantic alignment *C*_*XY*_ can be computed with respect to the semantic label *y*_*i*_’s for each task, shown in Figure 3C.

For the 3Dshape dataset, we take 4 latent visual features from each image, representing shape, scale, orientation and the color of the object. The remaining latent features (the wall’s and floor’s colors in the image) are fixed. Except for the shape feature, all the other latent visual features are continuous. Following the same procedure as for the Dsprites dataset, we select images that are either cube or cylinder for the shape feature. For the continuous features, we binarized them. We then randomly sampled different visual-to-semantic mappings to generate a family of tasks. The input-semantic alignment is computed with respect to these new semantic features and is shown in Figure 3C.

#### Transfer learning tasks

For the transfer learning tasks, during phase 1 training, we use tasks generated before (”Synthetic tasks for probe semantic representation”) with input-semantic alignment *<* 0.1. For these tasks, the HPC net can form an abstract representation in the PFC layer, but the no-HPC net cannot. During phase 2 training, we use the same set of stimuli as in phase 1. And we use the representation from the PFC layer obtained in phase 1 as input to a two-layer ReLU neural network, which we regard as the motor pathway to be learned during phase 2. The output of the new task in this phase depends only on the semantic features of these stimuli from phase 1.

Denote the semantic features of the stimuli from phase 1 as (*y*_1_, *y*_2_, …, *y*_*D*_). The 1st class of tasks is those in which the output of the new task depends only on a subset of the semantic features and is invariant to the remaining ones. In Figure 4B, we showed simulation results for two example tasks for *D* = 4, the majority and parity tasks. The 1D output for the majority task is given as *y*_*new*_ = sign(*y*_1_ + *y*_2_ + *y*_3_), where sign is the sign of the sum, which is +1 for a positive number and *−*1 for a negative number. For the parity task, the 1D output is given as *y*_*new*_ = *y*_1_*y*_2_*y*_3_, which also takes values from *±*1. In both (*D* = 4), we use stimuli with the semantic feature *y*_4_ = +1 to train the downstream two-layer motor pathway and test the network’s performance on stimuli with *y*_4_ = *−*1. Because *y*_4_ is the irrelevant feature in both tasks.

For the 2nd class of tasks, we divide the semantic features into two sets: stimulus features and rule features. The outputs of the new tasks depend on the stimulus features but are also modulated by the rule features. In Figure 4B, we showed simulation results for two example tasks for *D* = 4, the context-dependent majority-minority task and the context-dependent reversal task. The 1D output for the majority-minority task is *y*_*new*_ = *y*_4_sign(*y*_1_ + *y*_2_ + *y*_3_), where sign is the sign of the sum, which is +1 for a positive number and *−*1 for a negative number. For the context-dependent reversal task, the outputs are 3-dimensional and are given as *y*_*new,i*_ = *y*_*i*_*y*_4_, which also takes values from *±*1. In both cases, we think of *y*_4_ as the rule feature that determines whether the target action is flipped for the inputs. In both (*D* = 4), we randomly sample a fraction of semantic features and use the corresponding stimuli to train the downstream two-layer motor pathway and test the network’s performance on stimuli with the remaining semantic features.

#### Input-based generalization tasks

Similar to previous tasks, each stimulus has *D* binary semantic labels. The output for each stimulus in this task is assumed to be the same as its semantic labels. The stimulus input representation will be a distorted version of the semantic representation. Specifically, in the simulation, the number of semantic features is *D* = 4. If the semantic label of the stimulus is (*y*_1_, *y*_2_, *y*_3_, *y*_4_), then its input representation will be (*y*_1_*y*_4_, *y*_2_*y*_4_, *y*_3_*y*_4_, *y*_4_). Intuitively, half of the hypercube is flipped depending on the feature *y*_4_. Therefore, in order to perform the task, the network needs to flip back the labels for half of the hypercube vertices. During training, we randomly sample a subset of stimuli to train all three network models shown in Figure 5A. After training, the network is tested on the remaining stimuli to compute the generalization performance.

### Parallelism score and Cross-condition generalization performance

Given the neural representation in a specific layer of a network, since all the training data form 2^*D*^ distinct classes based on their semantic labels **l** = (*l*_1_, *l*_2_, .., *l*_*D*_), we define the prototype representation of each class as the mean firing rate vector of all the stimuli **r**^*i*^ belonging to this class,

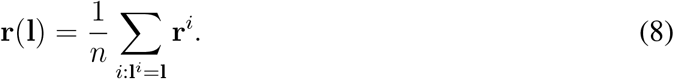

where **l**^*i*^ ∈ {*±*1}^*D*^ is the semantic labels for *i*the training stimulus.

To compute the parallelism score (PS), we examine how the neural representation changes when varying only the *k*th semantic label *y*_*k*_ while keeping other labels **l**_*\k*_ = *α* ∈ {*±*1}^*D*−1^ fixed,

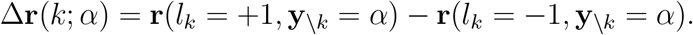

For different labels 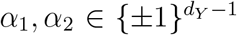, we quantify how consistent the direction of representation changes are when varying the *k*th label, using cosine similarity

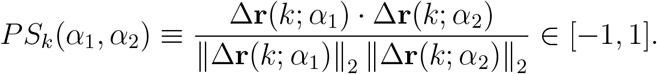

From this definition, if the representation change for the *k*th latent label is independent of other latent labels [Δ**r**(*k*; *α*_1_) = Δ**r**(*k*; *α*_2_)], then *PS*_*k*_(*α*_1_, *α*_2_) = 1.

The average parallelism score for the *k*th latent label is defined as the average cosine similarity for all distinct pairs (*α*_1_, *α*_2_),

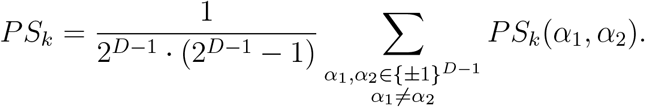

The overall *PS* for the neural representation is the average over all latent labels

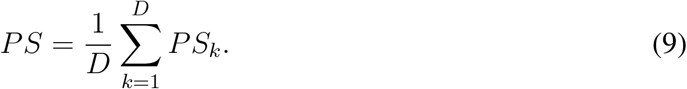

By the above definition, *PS* of a neural representation only depends on the inner product between the neural representation of all data pairs, **r**^*i*^ · **r**^*j*^. So we define the representation kernel matrix *K* ∈ ℝ^*P ×P*^ whose element is *K*_*ij*_ ≡ **r**^*i*^ · **r**^*j*^. This kernel matrix fully determines the *PS* for a neural representation.

In similar spirit, to compute the cross-condition generalization performance (CCGP) of the neural representation, we first fix two semantic labels *y*_*k*_ and *y*_*l*_. For all stimuli with *k*th semantic feature to be positive *y*_*k*_ = +1, we train a linear SVM decoder, using the representation of these stimuli as the input, to decode their *l*th semantic feature. After training, we test this SVM decoder on those stimuli with *k*th semantic feature to be negative *y*_*k*_ = *−*1. Denote this test performance as *CCGP* (*k, l*). This procedure can be done for all pairs of *k, l*, and we use the average over all pairs,

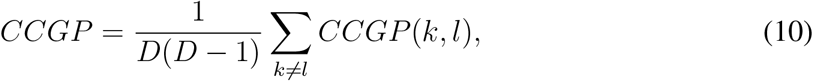

as the CCGP for this representation.

### Neural network models

Unless specified, all the network models are trained using stochastic gradient descent.

#### HPC-PFC networks

The feedforward HPC net is a multi-layered neural network: input layer, HPC layer, PFC layer and the readout. The HPC layer also projects to a reconstruction layer. The loss function of the HPC net is

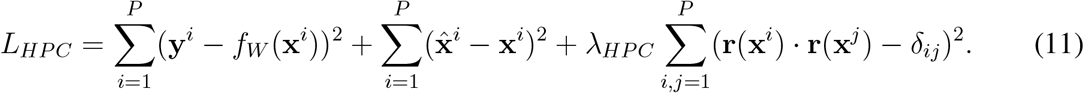

Here (**x**^*i*^, **y**^*i*^) is the input and output for the *i*th data point. *f*_*W*_ (**x**^*i*^) is the network output for *i*th data point. 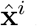 is the neural activity in the reconstruction layer. **r**(**x**^*i*^) is the neural activity for the *i*th data point in the HPC layer. The 1st MSE term depends on the weights in all layers of the network, except those in the HPC-to-reconstruction layer. The 2nd and 3rd term model the HPC module as an orthogonalized autoencoder and only depend on the input and reconstruction layer weights in the HPC module. During training, we introduced a small weight decay term and used a range of values for the regularization strength *λ*, and in all the plots, we show the one that gives the highest PS in PFC representation. The no-HPC net is a feedforward network with the same number of layers, with the same number of neurons in each layer, and its loss function only contains the 1st term in Eq.(11), which is a supervised loss.

In the HPC net with recurrent dynamics (Figure 3D), we use a vanilla RNN. Its loss is similar to Eq.(11) except that the 1st term is replaced by the MSE between RNN output and the actual output summed at the readout time points (2 time points in the simulation). A small weight decay is added during training. The corresponding no-HPC net has the same network architecture, except it does not have the reconstruction layer and only has the 1st term in Eq.(11) in its loss.

#### HPC-consolidation network model

In the HPC-consolidation net, there is one pathway (HPC-dependent pathway) that is the same as the HPC net before. Besides, there is another pathway from the input layer to the hidden layer and finally to the PFC (HPC-independent pathway). The total number of neurons in the HPC layer in the HPC-dependent pathway and the hidden layer in the HPC-independent pathway are matched to the total number of HPC neurons in the HPC net. During training, the HPC-dependent pathway is first trained in the same way as HPC net previously. After this training, the 2nd HPC-independent pathway (including the input-to-hidden weights and hidden-to-PFC weights) is trained to output the PFC representation learned during the 1st phase training, which we regard as the consolidation process. When computing test performance, we use the final network output only from the HPC-independent pathway.

### Analysis of optimal neural representation

#### Optimal representation only depends on the input-output kernel

We first show that the optimal representation of the HPC net depends only on the input and output kernels *K*_*X*_ and *K*_*Y*_. Therefore, the specific choices of the semantic-related readouts don’t matter as we discuss in the main text and in the previous section (”Synthetic tasks for probe semantic representation”). We introduce the input and output data matrix *X* = (**x**^1^, **x**^2^, …, **x**^*P*^) and *Y* = (**y**^1^, **y**^2^, …, **y**^*P*^). Also, we denote the input-to-HPC weights and bias as (*W*_1_, **b**_1_), HPC-to-reconstruction layer weights as *W*_1*d*_, the map from HPC’s preactivation to PFC as 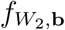, with the corresponding weights and bias (*W*_2_, **b**_2_) and the weights and bias of the output layer as (*W*_3_, **b**_3_)

The three terms in the loss of the HPC net can be written as

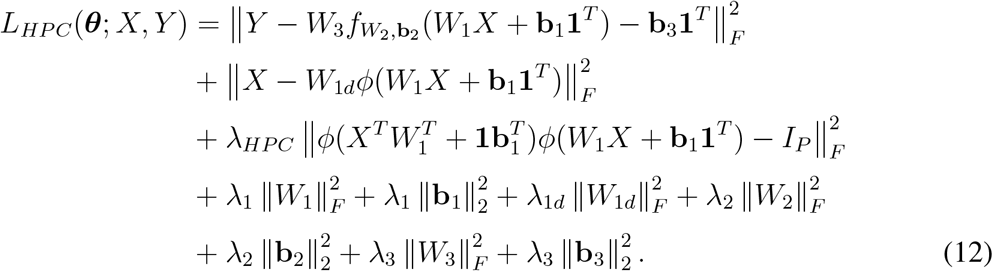

Here, we added the *l*^2^-regularization term on the weights and bias to account for weight decay. The trainable parameters ***θ*** = (*W*_1_, **b**_1_, *W*_*d*_, *W*_2_, **b**_2_, *W*_3_, **b**_3_) and we are interested in the optimal parameters argmin*L*(***θ***; *X, Y*).

Suppose we have two training dataset (*X, Y*) and (*X*′, *Y* ′), which share the same input and output kernel matrices

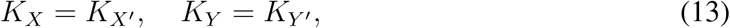

where *K*_*X*_ = *X*^*T*^ *X* and *K*_*Y*_ = *Y* ^*T*^ *Y*. Taking the singular value decomposition of the input and output data matrix, *X* = *UDV* ^*T*^, *X*′ = *U*′*D*′*V* ^*′T*^, *Y* = *O*Σ*W* ^*T*^, and *Y* ′ = *O*′Σ′*W*′, substituting in Eq.(13), we get

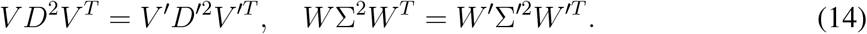

By the uniqueness of the singular value decomposition, we get *V* = *V* ′, *D* = *D*′, *W* = *W*′, and Σ = Σ′. So this means the two data matrices only differ by a rotation matrix

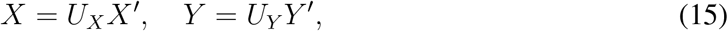

where *U*_*X*_ and *U*_*Y*_ are orthogonal matrices.

From Eq.(11), we notice that a rotation on the input data matrix is “transferred” to a rotation in the weight matrix *W*_1_. Therefore, we have

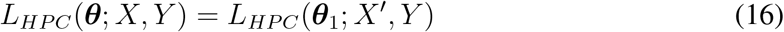

where ***θ***_1_ = (*W*_1_*U*_*X*_, **b**_1_, *W*_*d*_, *W*_2_, **b**_2_, *W*_3_, **b**_3_).

Next, we notice that only the 1st term in Eq.(11) depends on the output. Moreover, a rotation of the output data matrix can be transformed into a joint rotation in (*W*_3_, **b**_3_),

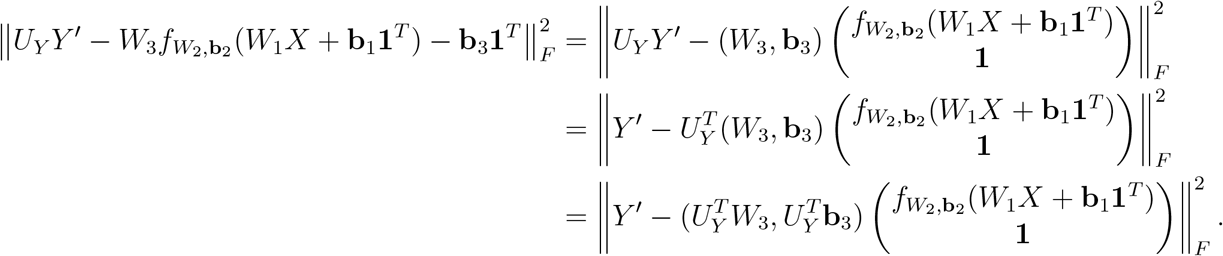

Therefore, this shows that

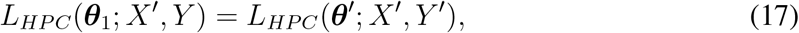

where 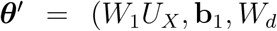, 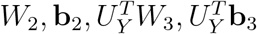). Combining the two results, we get *L*_*HPC*_(***θ***; *X, Y*) = *L*_*HPC*_(***θ***′; *X*′, *Y* ′). So if 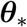 is a global minimum of 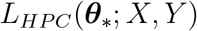, then 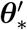 is a global minimum of 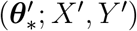.

Finally, we show that the network with parameter ***θ***_*\**_ and training data (*X, Y*) has the same representation kernel matrix as the network with parameter 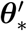 and training data (*X*′, *Y* ′). The representation kernel in the HPC layer for network with parameter ***θ***_*\**_ is

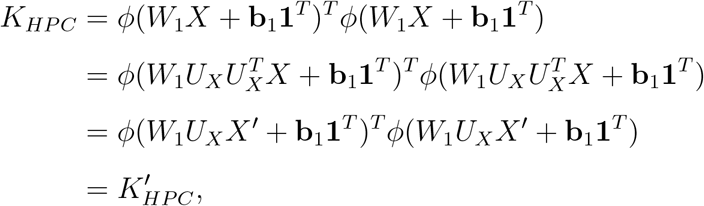

where 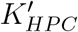 is the representation kernel matrix for the parameter 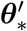.

Similar argument also applies to the PFC layer, whose representation kernel is

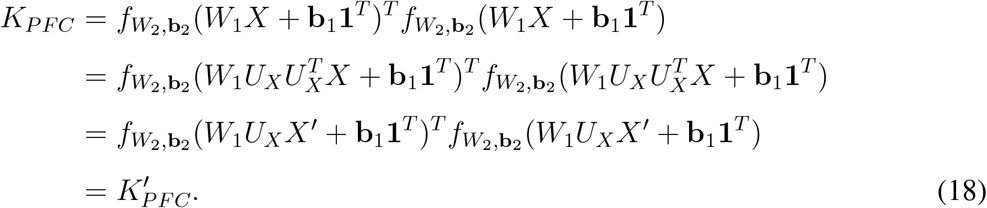

Therefore, the neural representation in the HPC and PFC layers at the global minima of *L*(***θ***; *X, Y*) are the same as the ones for *L*(***θ***′; *X*′, *Y* ′), as long as (*X, Y*) and (*X*′, *Y* ′) have the same input and output kernel matrices. While these analysis is done for HPC net, the one for no-HPC net is exactly the same without the last two terms in the loss [Eq.(11)].

#### Optimal representation for the downstream two-layer network

We consider the optimal PFC representation in the HPC net when its HPC representation is fixed. Specifically, we denote the neural activity matrix in the HPC layer for the training data as 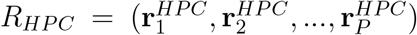. From the loss expression [Eq.(11)], the optimal PFC representation for a fixed HPC representation is determined only by the 1st term,

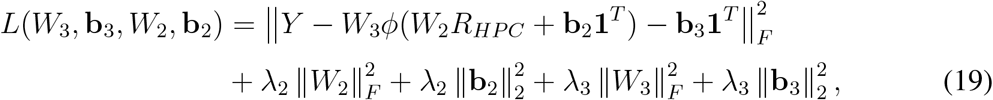

where *ϕ* is the element-wise ReLU function.

For fixed (*W*_2_, **b**_2_), we can minimize (*W*_3_, **b**_3_) to get

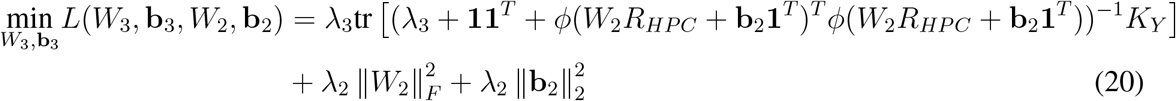

We make change of variables *H* ≡ *W*_2_*R*_*HPC*_ + **b**_2_**1**^*T*^ ∈ ℝ^*N ×P*^, where *N* is the number of neurons in PFC layer and *P* is the number of training data points. The above objective becomes a function of *H*,

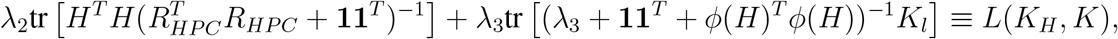

where *K*_*H*_ ≡ *H*^*T*^ *H* and *K* ≡ *ϕ*(*H*)^*T*^ *ϕ*(*H*). We define the following space

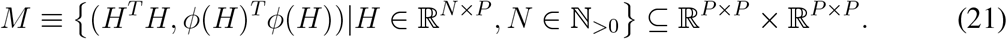

Since in the above objective, the 1st term is a linear function of *K*_*H*_ and 2nd term is a convex function of *K*, the objective is a convex function of (*K*_*H*_, *K*) over *M*.

Assume the HPC representation is orthogonal or more aligned with the representation kernel in the semantic space *K*_*l*_ = *K*_*Y*_,

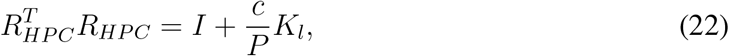

for *c* ∈ [0, *P*). We further assume that each semantic class has only one training data, so the total number of training data 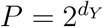.

Now we prove if *λ*_2_*λ*_3_ ≪ 1, the optimal solution will have the form

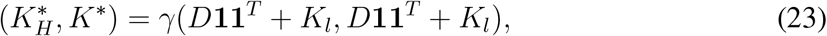

where 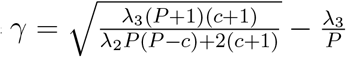.

To show this, for any (*K*_*H*_, *K*) = (*H*^*T*^ *H, ϕ*(*H*)^*T*^ *ϕ*(*H*)) ∈ *M*, we look at the loss value at the interpolating solution 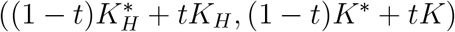 as

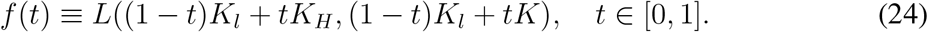

Since *f* (*t*) is composed of a linear function and a convex function, it is a convex function. We will prove that *f* is minimized at *t* = 0. The 1st derivative of *f* at *t* = 0^+^ is

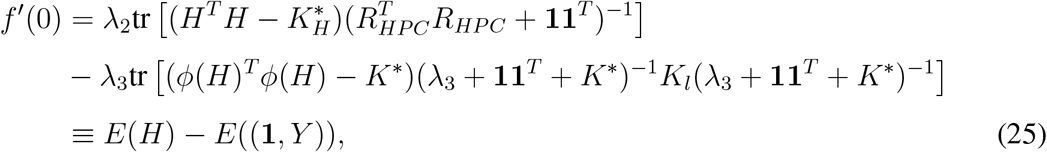

where the function *E*(*A*) for *A* = (**a**_1_, **a**_2_, **a**_3_, ..**a**_*N*_)^*T*^ ∈ ℝ^*N ×P*^ is defined as

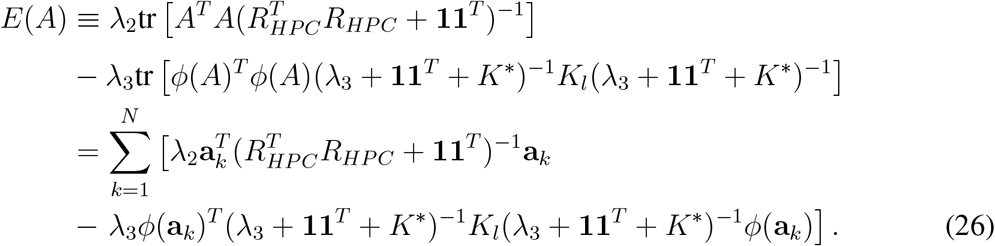

Using the definition of *K*_*l*_, *K*^*\**^, 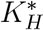 and *Y*, it can be checked that

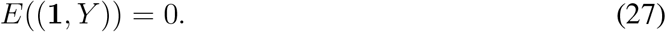

On the other hand, we will show that *E*(*H*) *≥* 0, ∀*H*. For the matrix *H* = (**h**_1_, **h**_2_, **h**_3_, ..**h**_*N*_)^*T*^ ∈ ℝ^*N ×P*^ Each term in the summation of Eq. (26) has the form

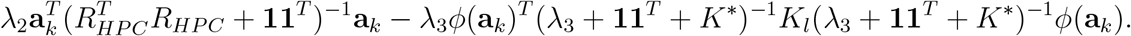

The 1st term can be lower bounded by

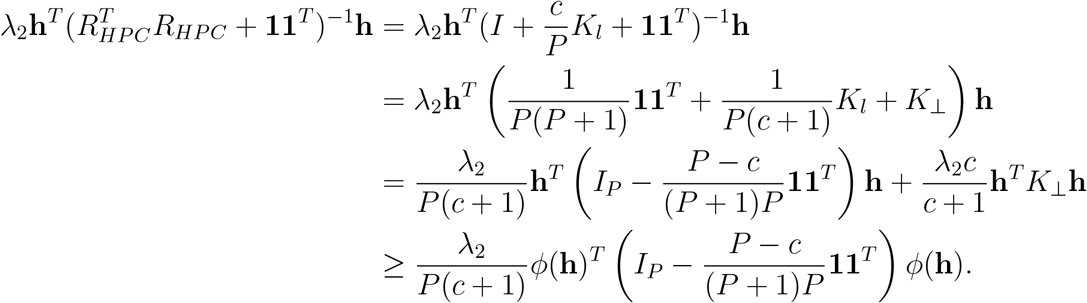

Here we use the fact that *ϕ* is ReLU. *K*_⊥_ is the projection matrix to the orthogonal subspace of the column space of *K*_*Y*_, (Range*K*_*Y*_)^⊥^.

Combining this with the 2nd term and using the definition of *K*^*\**^ and 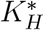, we get

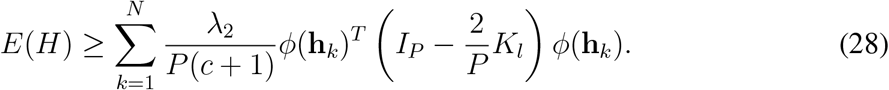

By definition, the representation kernel in the semantic space is generated by labels on *D*-dimensional hypercube, *K*_*l*_ = *L*^*T*^ *L*, where the 2^*D*^ column vectors of *L* are the vertices of the hypercube. This matrix *L* is known to have the following isotropy property [65],

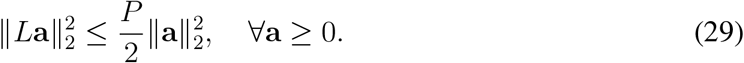

Applying this to the above lower bound, we get

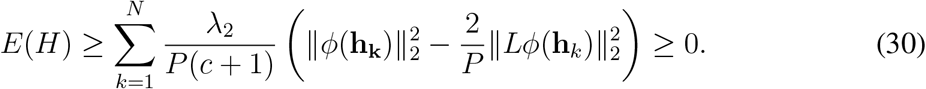

Altogether, we get

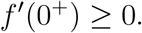

When equality attains [65], it requires that all row vectors **h**_*k*_ are selected from the set of vectors 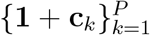 where **c**_*k*_’s are the row vectors of *L*. **c**_*k*_’s are also the eigenvectors of *K*_*l*_ [65]. In other word, if any of the row vectors is not from this subset, the strict inequality holds. Since *f* (*t*) is a convex function on [0, 1], *f*′(*t*) is monotonically increasing. This implies it attains the minimum at the left end-point *t* = 0.

In the case when equality holds, we need to check the 2nd-order derivative at *t* = 0^+^

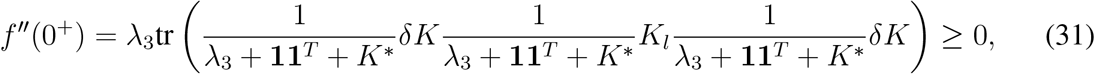

where *δK* ≡ *ϕ*(*H*)^*T*^ *ϕ*(*H*) *− K*^*\**^. Since the row vectors of *H* are linear combinations of **1** and **c**_*k*_, *δK* has the form 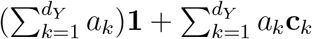. So the 2nd derivative vanishes if and only if

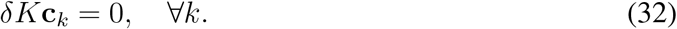

This implies *δK* = 0. In other word, if *δK* is nonzero, *f*^*′′*^(0^+^) *>* 0 and since the function is convex on [0, 1], it is minimized at *t* = 0.

So in summary, we showed that *f* (*t*) is minimized at *t* = 0. Therefore,

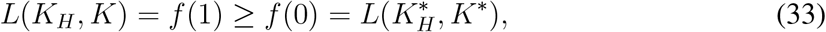

for any (*K*_*H*_, *K*) ∈ *M*. Equality holds only if *K* = *K*^*\**^ ∝ *d*_*Y*_ **11**^*T*^ + *K*_*l*_, which corresponds to the representation geometry in the semantic space.

## Data and code availability

No new experimental data was collected in this study. Source Data files are provided for all figures. Computer code to reproduce model simulations is available in the Github repository: https://github.com/BinW3233/AbstractRep.git.

**Figure S1:**
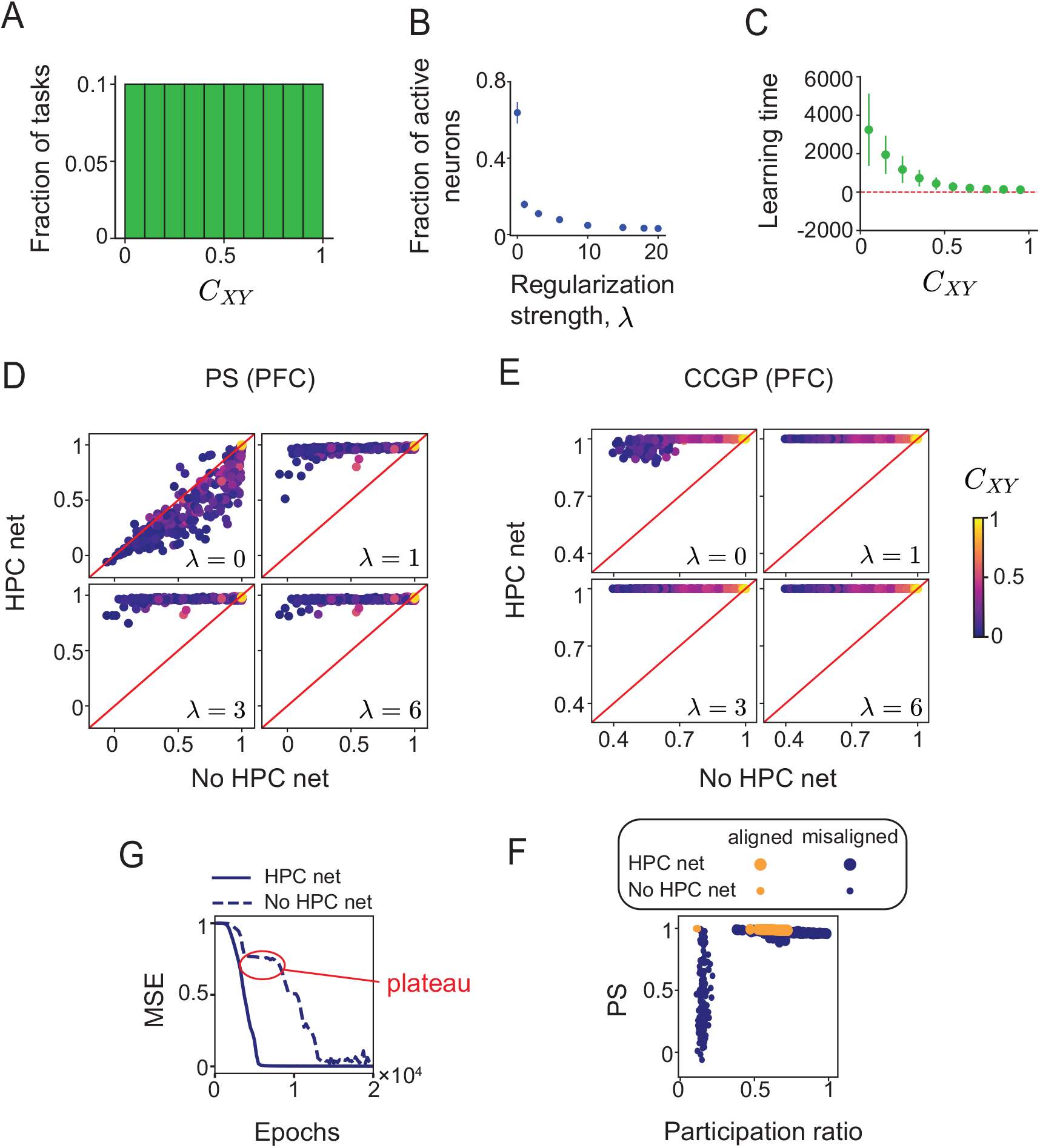
(A) The input-semantic alignment values are uniformly distributed across tasks. (B) Average learning time difference between the HPC net and the No HPC net. HPC net learns faster across different input-semantic alignment values. (C) Increasing the regularization strength automatically sparsifies the hidden representation of HPC. (D,E) The advantages of PFC in learning higher PS and CCGP persist across different values of *λ >* 0. (G) For mis-aligned tasks, the learning curve of the No HPC net shows a longer plateau than that of the HPC net. (F) When forming an abstract representation, increasing the dimensionality of the HPC representation has a larger effect on misaligned tasks than on aligned ones.

**Figure S2:**
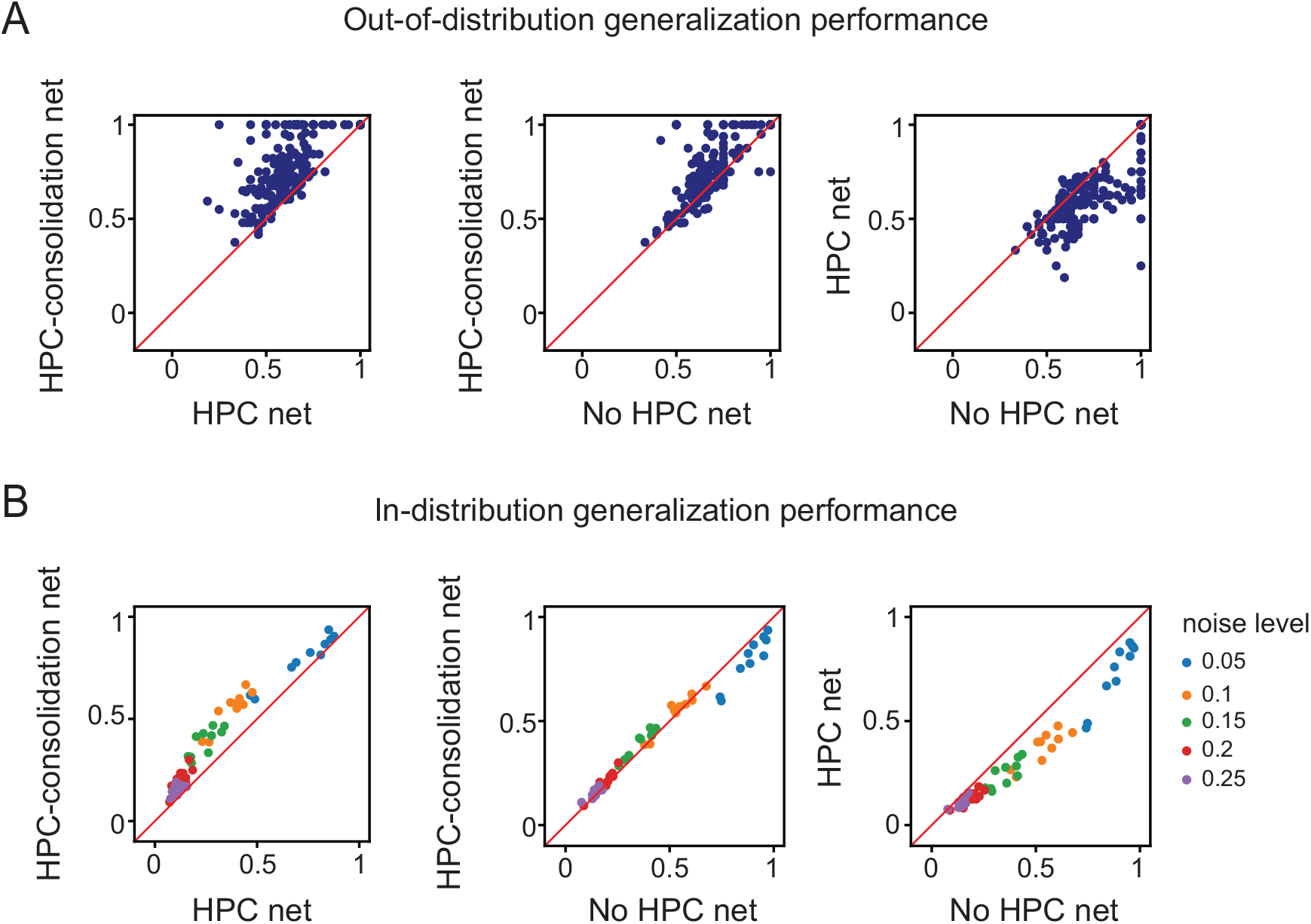
(A) Comparison of out-of-distribution generalization performance in the three network models. HPC-consolidation net consistently outperforms the other two network models. (B) Comparison of in-distribution generalization performance in the three network models. No single model consistently outperforms the others.

**Figure S3:**
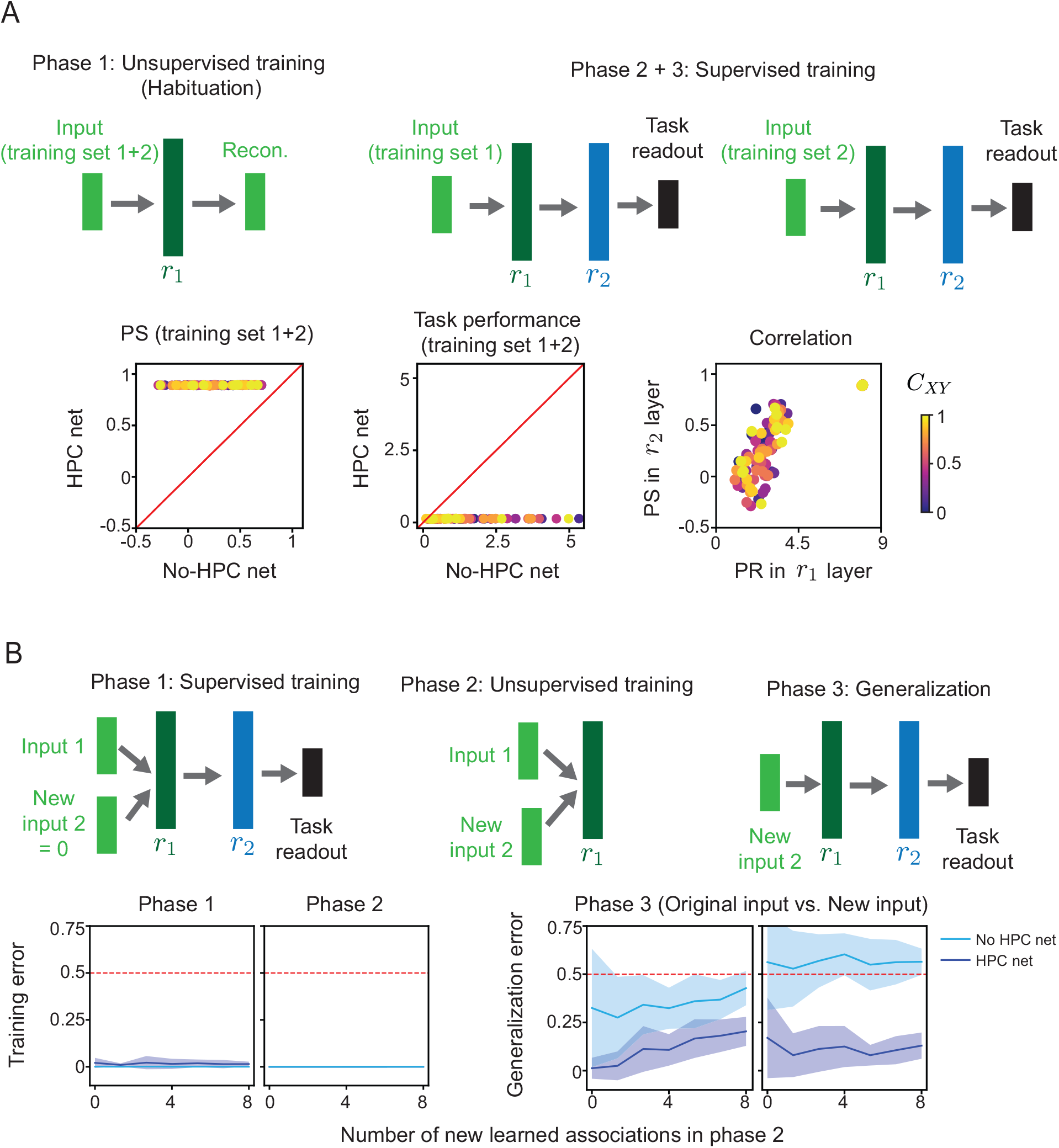
(A) High-dimensional HPC representation reduces interference during task learning and support abstract representation during sequential learning settings. (Top) The continual learning setting has three phases. During the 1st phase, the HPC layer (and the corresponding layer in No-HPC net) is trained in an unsupervised way to reconstruct the inputs. This phase models the process of Habituation where the animals get familiar with the stimuli. The difference between the HPC and No-HPC net is this phase is whether there is an orthogonalization regularization or not. During the 2nd and 3rd phase, both networks are trained as in the main text. The difference is that only half of the training data is used during each phase. After the three phase training, the PFC representation in the HPC net has a higher PS than the No-HPC net. HPC net has much better performance than the No-HPC net on all the training data. There is a positive correlation between the dimensionality in *r*_1_ layer and the abstractness of the representation in *r*_2_ layer. (B) High-dimensional HPC representation enables rapid generalization through orthogonalized binding mechanism. (Top) Three phases of the process: in phase 1, The network is trained in a supervised way, similar to the setup in Figure 2. After phase 1 training, in phase 2, we train both HPC net and no-HPC net unsupervisedly as autoencoders where we, for each stimulus input in phase 1, associate a new stimulus input to it. The idea is that the network will learn to “bind” the original stimulus input to the new input. The difference between the two networks is whether there is the orthogonal regularization in the objective function or not. In phase 3, we give the new inputs used for associative learning in phase 2 to the network and test whether the network gives the correct output (which is the target output used in phase 1). (Down) Shown on the left are the training errors during phase 1 and phase 2. both networks learn perfectly on the training data. In phase 3, the error in the HPC net is much smaller than in no-HPC net both when using the original input in phase 1 or the new input used in phase 2. This suggests that having the orthogonal regularization helps to bind new inputs to the old ones with less interference.

## Acknowledgments

The authors thank Matteo Alleman, Qihong(Q) Lu and other members in Fusi lab for their feedback and comments throughout the project. We are grateful to Arianna Pepin and Drew J Baughman for administrative support. This work is supported by NSF DBI-2229929 (ARNI), NIH NINDS K99NS138578 (WJJ), the Simons Foundation (542983SPI, SF), the Swartz Foundation, the Gatsby Charitable Foundation, and the Kavli Foundation.

## Competing interests statement

The authors declare no competing interests.

## Notes

### Competing Interest Statement

The authors have declared no competing interest.

